# Deep sequencing of High Plains wheat mosaic virus from sweet corn to guide seed health testing reveals multiple variants for all eight genome segments and two major isolate types

**DOI:** 10.64898/2026.08.25.746265

**Authors:** Jennifer R. Wilson, Erik W. Ohlson, Kristen J. Willie, Nitika Khatri, Lindsey J. du Toit

## Abstract

High Plains wheat mosaic virus (HPWMoV) is a wheat and maize-infecting virus of phytosanitary concern due to its potential for seed transmission. Recent phytosanitary restrictions have required sweet corn seed lots to test negative for HPWMoV prior to import into certain countries. To inform the design of more sensitive and broad-spectrum diagnostic primers for seed health testing and phytosanitary certification, we performed deep sequencing of HPWMoV-positive tissue collected from fields in two major sweet corn seed production regions in the Pacific Northwest, the Columbia Basin and Treasure Valley. Virus-like particle enrichment prior to Illumina sequencing facilitated near complete genome coverage (>95%) for the 21 HPWMoV isolates sequenced. *De novo* assembly of the eight viral genome segments revealed high levels of diversity for each segment, with at least two variants identified for each RNA and three variants for RNA3, RNA6, and RNA8. Within each sample, only one variant per RNA segment was usually present, with the notable exception of RNA3, sorting each isolate into what we designated type A and type B isolates. All but one previously sequenced HPWMoV isolate can be sorted into these two types. Two samples contained at least two variants for every RNA, totaling 17 genome segments, potentially representing a co-infection of type A and type B isolates. Despite this variability, we successfully designed two primer and probe sets for reverse transcription-quantitative polymerase chain reactions (RT-qPCR) that detected all 20 isolates tested in a duplex diagnostic assay, making the assay suitable for seed health testing for HPWMoV.

## 1. Introduction

High Plains wheat mosaic virus (HPWMoV) is an eriophyid mite-transmitted virus that infects corn and wheat as well as some other poaceous crops and grasses (Seifers et al., 1998). HPWMoV belongs to the species *Emaravirus tritici* in the family *Fimoviridae*. Other previous common names for HPWMoV in the literature include High Plains virus (HPV), wheat mosaic virus (WMoV), and maize red stripe virus (MRSV/MRStV). HPWMoV can cause mosaic and chlorotic or red streaking symptoms on sweet corn leaves, sometimes leading to necrosis and even plant death if plants are infected at an early growth stage. HPWMoV is transmitted by the wheat curl mite, *Aceria tosichella*. Due to the potential for seed infestation and transmission (Forster et al., 2001; Nischwitz, 2020), the virus has recently been of phytosanitary concern. Restrictions were put in place by some countries requiring sweet corn seed lots to be certified “free” of HPWMoV prior to import from the United States (*Establishes emergency measure to prevent the entry of High Plains virus (HPV) and wheat streak mosaic virus (WSMV) in corn seed from all origins*, 25 Nov 2022; *Import Health Standard: Seeds for Sowing*, Aug 30 2024).

The most common method of phytosanitary certification is seed health testing, increasingly by reverse transcriptase-polymerase chain reaction (RT-PCR) assays. However, the lack of RNA sequence information for HPWMoV, diversity among the isolates sequenced to date, and complexity of the genome have hindered effective diagnostic primer design. HPWMoV has a negative-sense RNA genome with eight genome segments, but two versions of RNA3 are often present, designated RNA3A and RNA3B (Tatineni et al., 2014). The primers currently used for seed health testing (Arif et al., 2014; Lebas et al., 2005) were developed prior to the first full genome of HPWMoV being sequenced, and both primer sets target RNA3, which is a poor target for diagnostics due to its heterogeneity. Indeed, both primer sets match the RNA3B reference sequence but have several mismatches with the RNA3A sequence. As many as 17 other HPWMoV isolates have been sequenced since (Albrecht et al., 2022; Candresse et al., 2026; Hodge et al., 2020; Jones et al., 2023; Stewart, 2016), but due to the difficulty of obtaining the full-length genome without virus enrichment, most of these sequences are partial, with only six full-length HPWMoV genome sequences available prior to this study.

There is specifically a lack of sequence information for isolates collected from corn, with only one isolate partially sequenced to date (Stewart, 2016), and none sequenced from sweet corn. Geographically, only one isolate has been sequenced from the Pacific Northwest, where two of the major sweet corn seed production regions are located in the United States: namely, the Columbia Basin along the Columbia River in central Washington and northcentral Oregon, and the Treasure Valley surrounding the Snake River in southwest Idaho and eastern Oregon. The previously sequenced isolate was collected from a wheat plant in Idaho, but not from the Treasure Valley (Hodge et al., 2020).

To improve primer design for seed health testing for HPWMoV for phytosanitary certification, we sequenced HPWMoV isolates collected from sweet corn plants in the Columbia Basin and Treasure Valley. We report the full genome sequences of 21 new HPWMoV isolates, greatly expanding our knowledge of HPWMoV genome diversity, as well as the development of two new diagnostic primer and probe sets with broad recognition of HPWMoV isolates that could be used for seed health testing.

## 2. Material and Methods

### 2.1 Virus isolate collection

Sweet corn plants with HPWMoV-like symptoms were sampled and shipped to the USDA-ARS Corn, Soybean & Wheat Quality Research Unit in Wooster, Ohio. Leaf samples collected from the same field were considered the same isolate. The number of plants sampled was not always recorded but samples known to be collected from a single plant are indicated in **Table 1**. Additional metadata for each sample is presented in **Table 1**, and collection locations are noted in **Figure 1**. All isolates were designated with the last two digits of the year of collection, the two-letter abbreviation for the state, and a number (e.g. 15WA1, 22OH1, 23ID3, etc.). For samples collected in 2022, total nucleic acid was extracted using grape extraction buffer (1:20 w/v; GrEB: 0.05 M sodium carbonate buffer pH 9.6, 2% polyvinylpyrrolidone-40, 0.2% bovine serum albumin, 0.05% Tween-20) and denatured in GES buffer (GES: 0.1 M glycine-NaOH pH 0.0, 50 mM NaCl, 1 mM EDTA, 0.5% Triton X-100, and 10 mM DTT). For samples collected in 2023, RNA was isolated from leaf samples using the Direct-zol RNA Miniprep Plus kit (Zymo Research) following the manufacturer’s instructions. All samples were tested for the presence of HPWMoV and WSMV using a one-step reverse transcription-polymerase chain reaction (RT-PCR) using Superscript III (ThermoFisher) for RT and Green GoTaq Polymerase (Promega) for PCR, as previously described (Xie et al., 2021). Two primer pairs were used for detection of HPWMoV: one targeting RNA3 (Lebas et al., 2005) and another targeting RNA2 (**Supplementary Table 1**). One primer set was used for detection of WSMV (Byamukama et al., 2016). Samples testing positive for HPWMoV were stored at −80°C, and 0.5 g of tissue was used for propagation via vascular puncture inoculation (VPI) of the susceptible field corn inbred Oh28, as previously described, using Buffer B from Louie et al. (2006) which is comprised of 0.05 M sodium phosphate, 0.005 M ethylenediaminetetraacetic acid (EDTA), and 0.01 M sodium sulfite buffer, at pH 7.8. VPI propagation also confirms the presence or absence of WSMV which can also be transmitted via VPI, but at a higher rate than HPWMoV.

**Figure 1.**
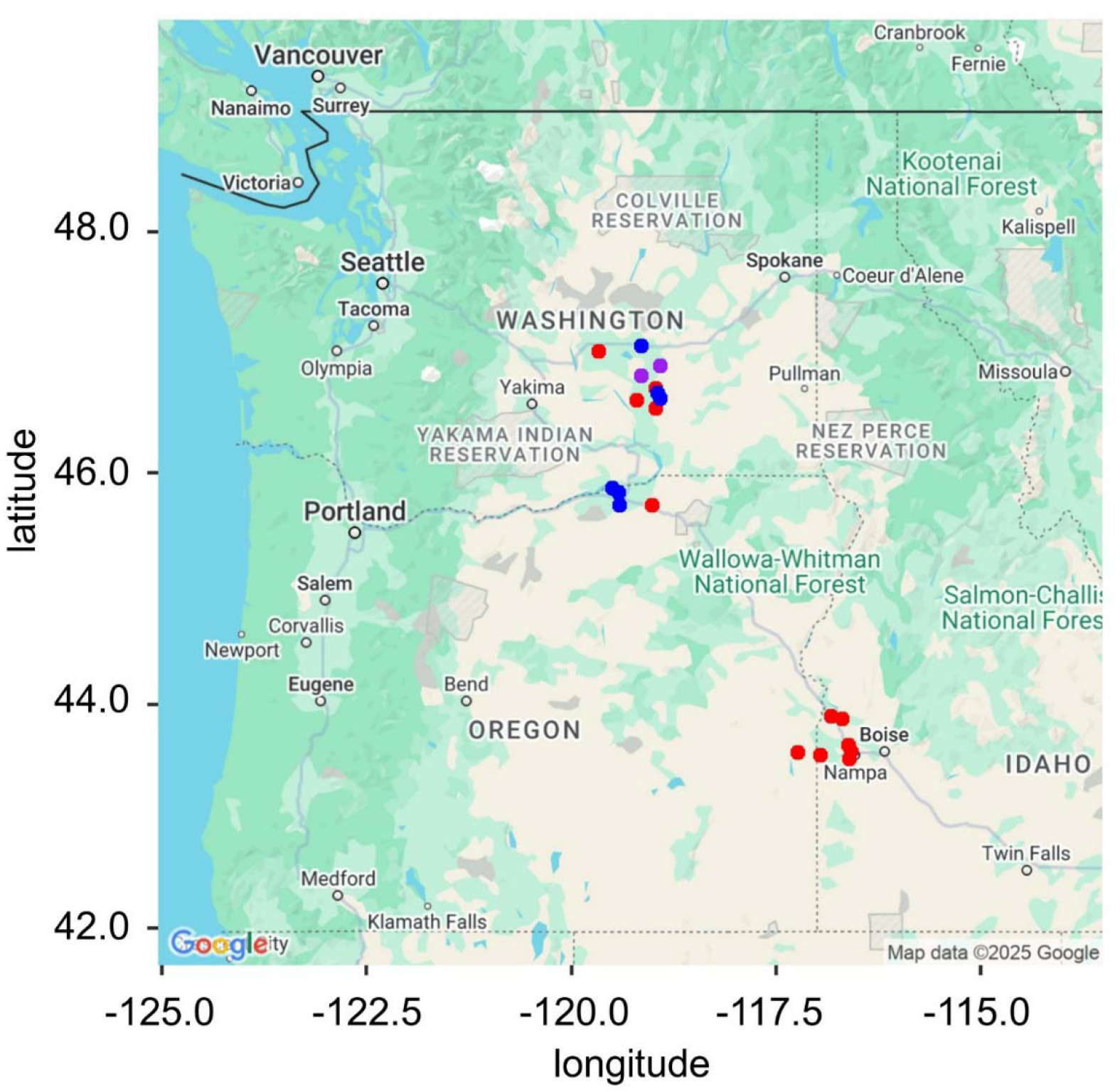
Samples collected from the two major sweet corn seed production regions in the Pacific Northwest USA. Dots indicate the approximate location of collection of 20 samples from Washington, Oregon, and Idaho. Dots are colored according to the type of HPWMoV isolates sequenced from that sample: red indicates an A-type isolate, blue indicates a B-type isolate, and purple indicates that both A and B isolates types were recovered from the sample. GPS coordinates of nearby towns were used when exact field GPS coordinates were not collected. All points have been “jittered” to protect the exact location of sampled farms.

**Table 1.** Metadata associated with the 21 HPWMoV isolates collected and sequenced.

| **Sample** | **Year** | **Collection Location** | **Germplasm<sup>a</sup>** | **WSMV<sup>b</sup>** |
| --- | --- | --- | --- | --- |
| 15WA1 | 2015 | WA | (not recorded) | - |
| 22OH1 | 2022 | Wayne County, OH | fresh market hybrid | - |
| 22ID1 | 2022 | Canyon County, ID | inbred | - |
| 22WA1 | 2022 | Grant County, WA | processing hybrid | + |
| 22WA2<sup>c</sup> | 2022 | Adams County, WA | inbred | - |
| 23ID1 | 2023 | Canyon County, ID | inbred | + |
| 23ID2 | 2023 | Canyon County, ID | inbred | - |
| 23ID4 | 2023 | Canyon County, ID | inbred | - |
| 23ID5 | 2023 | Canyon County, ID | inbred | - |
| 23ID6<sup>c</sup> | 2023 | Canyon County, ID | inbred | - |
| 23OR1 | 2023 | Morrow County, OR | processing hybrid | + |
| 23OR2 | 2023 | Morrow County, OR | processing hybrid | + |
| 23OR3 | 2023 | Morrow County, OR | processing hybrid | - |
| 23OR4 | 2023 | Malheur County, OR | inbred | - |
| 23OR5 | 2023 | Umatilla County, OR | processing hybrid | - |
| 23WA1<sup>c</sup> | 2023 | Adams County, WA | processing hybrid | - |
| 23WA2<sup>c</sup> | 2023 | Adams County, WA | processing hybrid | - |
| 23WA3 | 2023 | Franklin County, WA | inbred | - |
| 23WA4 | 2023 | Franklin County, WA | inbred | - |
| 23WA5 | 2023 | Adams County, WA | processing hybrid | - |
| 23WA6 | 2023 | Adams County, WA | processing hybrid | - |

<sup>a</sup> All samples were collected from sweet corn. Sweet corn plants were either hybrids for fresh market production, hybrids for canning and processing, or inbred parents for hybrid seed production.
<sup>b</sup> A subset of field collected samples was co-infected with wheat streak mosaic virus (WSMV) detected via RT-PCR and confirmed during isolate propagation
<sup>c</sup> This sample was recorded to be collected from a single plant.

### 2.2 Virus-like particle enrichment and RNA isolation

Original field-collected tissue was used, when possible, if a sufficient amount of high-quality tissue was available. Else, wheat tissue infected via wheat curl mites (sample 22OH1) or maize tissue infected via VPI (samples 22ID1, 22WA1, 22WA2) was used (Table 2). Virus-like particles were isolated using a protocol adapted from Tatineni et al. (2009). Approximately 2.5 g of tissue was homogenized with a mortar and pestle in 1:4 w/v sodium citrate buffer (0.1 M sodium citrate, pH 6.5) with 0.1% β-mercaptoethanol. Homogenate was filtered through four layers of cheesecloth. Cell debris was pelleted by centrifugation for 10 minutes at 7,900 rpm (7,700 x *g*) in a Sorval SM24 rotor. The supernatant was collected, combined with 2% Triton X-100, and stirred for 10 minutes. Then, the extract was layered onto a 20% sucrose cushion and ultracentrifuged for 1.5 hours at 32,700 rpm (118,000 x *g*) in a Beckman Type 42.1 rotor. The resulting pellet was resuspended in sodium citrate buffer and centrifuged for 10 minutes at 7,900 rpm (7,700 x *g*) in the SM24 rotor. The supernatant was layered upon a second 20% sucrose cushion and ultracentrifuged for 1.5 hours at 35,500 rpm (139,000 x *g*) in the Type 42.1 rotor. The pellet was resuspended in 200 uL of 0.1 M potassium phosphate buffer, pH 7.0. One milliliter of Trizol® was added to the virus preparation and total RNA extracted using the Directzol RNA Miniprep Plus kit (Zymo Research) following the manufacturer’s instructions.

**Table 2.**
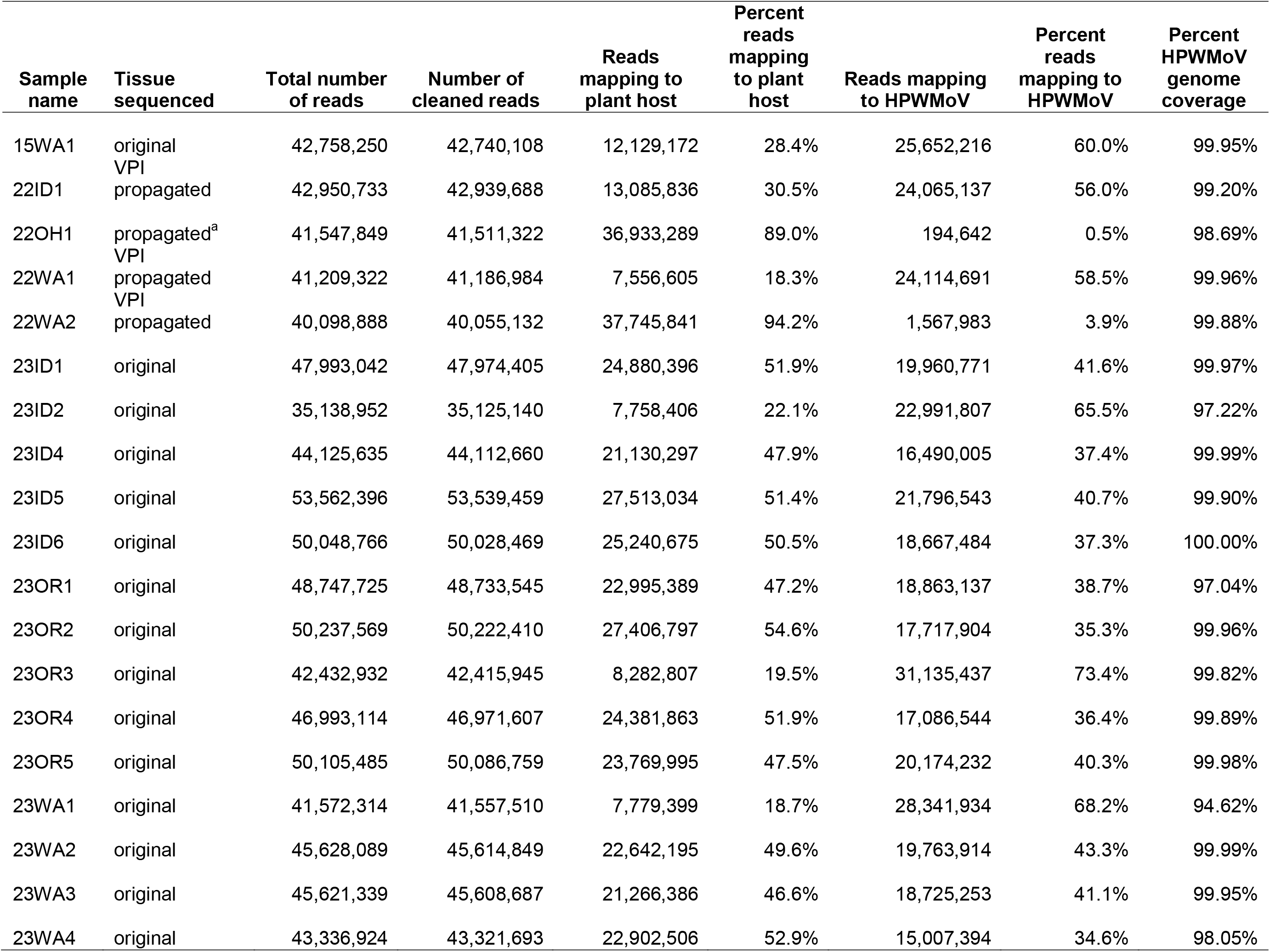
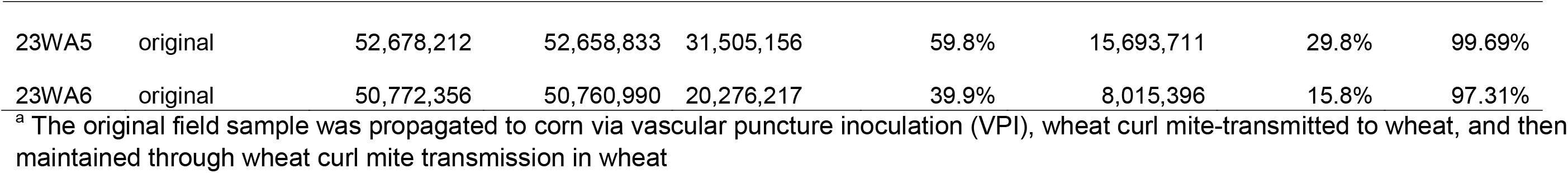
Summary of RNA sequencing results including host read subtraction and viral genome coverage for High Plains wheat mosaic virus (HPWMoV)

### 2.3 Library preparation and sequencing

Library preparation and sequencing was performed by the Genomics Shared Resource at The James Comprehensive Cancer Center at The Ohio State University. Ribosomal RNA depletion of total RNA was performed using the QIAseq FastSelect Plant rRNA removal kit (Qiagen) and total transcriptome libraries constructed using the NEBNext® Ultra II Directional RNA kit (New England Biolabs) following the manufacturer’s instructions. Libraries were sequenced with an Illumina Nova Seq X Plus with 100 bp paired end reads.

### 2.4 Bioinformatic analysis

Quality control (QC) of the demultiplexed paired end FASTQ files was performed using Trimmomatic v0.39 and default parameters (Bolger et al., 2014). Read quality was evaluated before and after QC using FastQC v0.12.1(Andrews, 2010). Subsequently, reads were mapped to the corresponding host corn (Zm-B73-REFERENCE-NAM-5.0) or wheat (IWGSC RefSeq v2.1) reference genome using the Burrows-Wheeler Aligner (BWA) 0.7.18 *mem* function (Li & Durbin, 2009).

Assembly was performed *de novo* using the - *rnaviral* function in SPAdes 3.15.5 (Prjibelski et al., 2020). The *de novo* scaffolds were quantified using Salmon v1.10.1 in mapping-based mode for each sample using k-mer length 31 (Patro et al., 2017). Scaffolds for which the transcripts per million (TPM) were less than 1,000 were excluded from further analysis. The *de novo* assembled scaffolds were mapped back to the reference HPWMoV genome assembly using minimap2 to determine RNA segment identity and to perform additional QC by correcting palindromic sequences and misassemblies (Li, 2018). Unmapped scaffolds were aligned to the HPWMoV reference genome using blastn within the BLAST+ software suite to identify more divergent sequences (Camacho et al., 2009).

Scaffolds that were palindromic or the result of concatenated RNA segments were manually corrected. Since HPWMoV begins and ends with identical 12-nt sequences, ends were manually trimmed to the reference genome, if necessary. Scaffolds representing alternative variants are reported when nucleotide identity was <95% between the primary and alternative variant(s), otherwise the full-length consensus sequence is reported with nucleotide ambiguity codes. For each isolate, RNA variants representing <5% of the total copies of that segment were excluded from subsequent analyses.

All phylogenetic trees and multiple sequence alignment figures were generated in Geneious Prime 2026.0.2. Phylogenetic trees were constructed using the neighbor-joining method and the Tamura-Nei genetic distance model with 100 bootstrap replicates. Multiple sequence alignments of isolates sequenced in this study were aligned using MUSCLE 5.1 and included as **Supplementary Data**. Consensus sequences were generated from a multiple sequence alignment of all sequences from this study of that variant (e.g. RNA1A) with a 90% cut-off for assignment of ambiguous bases. Multiple sequence alignments of isolates from previous studies were aligned to these consensus sequences using Clustal Omega 1.2.2. due to the large number of partial sequences. Genome segments from previously sequenced isolates were designated “A” or “B” based on their percent nucleotide similarity to the consensus sequences. Isolates were designated “type A” or “type B” based on the overall genome segment composition.

Recombination analysis was performed in RDP5 (Martin et al., 2021) on the HPWMoV sequences generated in this study, specifically the multiple sequence alignments included in the **Supplementary Data**. Recombination analysis was performed with all nine recombination detection algorithms (RDP, GENECONV, Bootscan, MaxChi, Chimaera, SiScan, 3Seq, LARD, and Phylpro) with default parameters (Gibbs et al., 2000; Holmes et al., 1999; Lam et al., 2018; Martin & Rybicki, 2000; Martin et al., 2005; Padidam et al., 1999; Posada & Crandall, 2001; Smith, 1992; Weiller, 1998). Recombination events are only reported if they were significant at p < 0.05 for at least seven of the nine prediction algorithms.

### 2.5 Sanger and Oxford Nanopore sequencing confirmation

Sanger sequencing was used to confirm the presence of both A and B variants for RNA1, RNA2, RNA4, and RNA7 in the samples 23WA1 and 23WA4. These RNA segments were selected for validation as typically only one variant of each of these RNAs was observed for all other samples. Primers specific to the A or B variant for RNA1, RNA2, RNA4, and RNA7 for 23WA1 and 23WA4 were designed (**Supplementary Table 1**), and RT-PCR performed as described above but using PrimeStarGXL as the DNA polymerase on the original RNA used for Illumina sequencing for 23WA1 and 23WA4. The resulting PCR products were purified using the Monarch Spin PCR & DNA Clean-Up Kit (New England Biolabs) and Sanger sequenced with the same primers (Genewiz). The presence of all three variants of RNA3 were also confirmed in samples 23WA1 and 23WA4 in the same manner.

Oxford Nanopore (ONP) long-read sequencing was also used to assess four unique insertions >30 bp found in some RNA variants from samples 22WA1 (RNA3B), 23WA1 (RNA4A), and 23WA4 (RNA2B, RNA6B) and 2 insertions present only in 23OR2 and 23WA1 in each of RNA7B and RNA8B. ONP was also used to sequence regions with a predicted recombination event in RNA5 from isolates 23WA5 and 23WA6, and from RNA7 from isolates 23WA6 and 15WA1. PCR primers were designed that spanned each insertion or recombination event and RT-PCR was performed using PrimeStarGXL polymerase on the original RNA used for Illumina sequencing for the respective sample. PCR products were purified and sent for ONP sequencing (Plasmidsaurus Inc.). Sequences from the predicted recombination regions are included in the **Supplementary Data**. None of the insertions was validated and, therefore, the insertions were manually removed from the sequences.

### 2.6 Design and testing of new diagnostic primers

Prior to Illumina sequencing of the HPWMoV isolates, a primer and probe set suitable for reverse transcription quantitative PCR (RT-qPCR) was designed for RNA2 by generating a multiple sequence alignment of all RNA2 sequences available in GenBank at the time of access (17 May 2023). The IDT Primer Quest Tool was used to identify primer and probe sequences with desirable properties for RT-qPCR, such as melting temperature. Degenerate bases were used in the primers to capture all sequence variability, but no mismatches or degenerate bases were tolerated in the probe. After Illumina sequencing of the 21 new HPWMoV isolates, a new primer and probe set was designed to RNA4 using the same parameters and based off a multiple sequence alignment of all isolates sequenced in this study, plus the reference sequence. Primers and probe were synthesized by Eurofins with the RNA2 probe (5’-CTGATGGTGAAAGACAAATGTTTCCTG-3’) conjugated to Cy5® with TAO™ and Iowa Black® RQ quenchers. The flanking RNA2 primers are: HPV RNA2 869F (5’-ATCCTTGASTATGTGAGARCC-3’) and HPV RNA2 947R (5’-AAAGACARCCGAGCATACA-3’). The RNA4 probe (5’-TGACATGCATACAGGTAATTTGGCAAC-3’) was conjugated to 5’ 6-carboxyfluorescein (FAM) with ZEN™ and 3’ Iowa Black® FQ as quenchers. The flanking RNA4 primers are HPWMoV RNA4 941F (5’-CTGTKGCYTCACCTGGRATAG-3’) and HPWMoV RNA4 1031R (5’-TRGTCCAAATTGTRTCAGGNA-3’).

Original field tissue was used for RT-qPCR testing, where possible. Else, propagated virus-infected leaf tissue was used. Seeds from isolates 22WA2 and 23WA6 were collected from the same plant in the field and shipped along with the leaf tissue. Infested seeds from isolate 22OH1 were generated by mite inoculation of the sweet corn inbred ARZM 19 057, followed by transplanting to the field, self-pollination, and seed collection. Infested seeds for isolate 23ID2 were generated by VPI of the experimental field corn hybrid Wf9 x Oh51A, which were planted, grown to maturity, and self-pollinated in a BSL-2 containment greenhouse. All seeds were dried to <15.5% moisture content and stored at 4°C prior to testing. Seeds and leaf tissue for RT-qPCR testing were cryoground in liquid nitrogen with a mortar and pestle, and RNA was extracted using the Direct-zol RNA Miniprep Plus kit.

One-step RT-qPCR was performed using the TaqMan Fast Virus 1-Step Master Mix (Applied Biosystems) in a 20 uL reaction volume with 5 uL of RNA template,250 nM of RNA2 primers, 300 nM of RNA4 primers, and 200 nM of each probe. Each RT-qPCR was performed in a CFX96 C1000 thermal cycler (Bio-Rad) with the following parameters: RT for 5 minutes at 50°C, initial denaturation at 95°C for 2 seconds, 35 cycles of 95°C for 15 seconds, and 60°C for 60 seconds. The fluorescence threshold was manually set to be at the midpoint of the exponential amplification curve for each probe.

## 3. Results

### 3.1 Twenty HPWMoV-infected sweet corn samples were collected from the two major sweet corn seed production regions in the Pacific Northwest

Over the 2022 and 2023 growing season, 19 samples were collected from sweet corn in Washington, Oregon, and Idaho, with at least five samples collected from each state, covering both of the major sweet corn seed production regions for the USA: the Columbia Basin and the Treasure Valley (**Figure 1**). One additional sample was collected from Ohio in 2022.

All samples tested positive for HPWMoV via RT-PCR. Four samples also tested positive for WSMV and WSMV infection in these samples was confirmed during propagation via VPI (**Table 1**). Samples included an even distribution of sweet corn hybrids and inbreds. Inbreds were grown for seed production whereas most of the hybrids were from crops grown for processing, with the exception of the one sample from Ohio which was collected from a hybrid grown for fresh market production. A sample collected in Washington State in 2015 was also included in the sequencing, though collection location and germplasm information were not recorded for this sample.

### 3.2 Partial virus purification to enrich for viral sequences enabled near complete viral genome coverage

A partial virus purification was performed on each leaf sample to enrich for virus-like particles, followed by RNA extraction, rRNA depletion, library preparation, and Illumina sequencing. Where possible, original field tissue was used for the extraction but in some cases the virus had been propagated one or more times via vascular puncture inoculation (samples collected in 2022) or mite transmission (22OH1), as indicated in **Table 2**. All samples were sweet corn tissue, except for 22OH1 which was originally collected from sweet corn but maintained in wheat.

An average of 45 million reads were obtained for each sample (**Table 2**). After QC, on average, 46.3% of reads per sample mapped to the host plant genome (range of 18.3 to 94.2%) and 40.9% mapped to the HPWMoV genome (range of 0.5 to 73.4%), indicating that, in general, the partial virus purification procedure was effective at enriching for viral sequences relative to host-derived sequences. The sample with the fewest reads mapping to HPWMoV, with only 0.5% viral reads, was isolated from wheat rather than sweet corn. HPWMoV has been successfully sequenced from wheat before, so the low proportion of viral reads may be due to low levels of infection in the wheat plants that were sampled. For the two other samples with the next lowest proportion of viral reads, 22WA2 rapidly caused plant death when inoculated via VPI, and 23WA6 was from poor quality tissue that had begun to decay by the time the sample was received. In both cases, fewer living plant cells were present, which are needed to isolate intact virions.

Considering the high number of reads, the relatively short viral genome, and successful enrichment of viral sequences, an extremely high coverage depth was obtained for the HPWMoV genome for all samples, with an average depth of 210,000x. Even the sample with the fewest viral reads, 22OH1, had an average sequencing depth of 1,930x across the HPWMoV genome. Near complete coverage of the HPWMoV genome was obtained for all samples with average genome coverage of 99.1% (range of 94.62 to 100%, **Table 2**).

### 3.3 *De novo* assembly reveals two or three major variants for each RNA segment

Considering the previous finding of multiple variants of RNA3 (Tatineni et al., 2014), in addition to map-based assembly, *de novo* assembly of sequencing reads was conducted to identify novel variants of each RNA segment. *De novo* assemblies were then mapped back to the HPWMoV reference genome to determine RNA segment identity.

When phylogenetic analysis was conducted on all sequences for each RNA segment, it became apparent that the sequences largely separated into two or three distinct clades, as demonstrated by RNA2 and RNA3, respectively (**Figure 2**). For RNA3, the RNA3A sequence from the reference isolate determined by Tatineni et al. (2014) clustered with one group of 15 RNA3 variants, which have also been designated “RNA3A”. The RNA3B reference sequence clusters with 15 other RNA3 sequences from this study, which have also been designated “RNA3B”. A third clade of 8 RNA3 sequences separated from RNA3A and RNA3B, which were designated “RNA3C” (**Figure 2B**). In contrast, RNA2 sorted into two major clades: 15 sequences that grouped with the RNA2 reference sequence, deemed “RNA2A”, and 8 sequences that formed a separate clade, designated “RNA2B” (**Figure 2A**). In a similar manner, all clades of sequences for each RNA segment were assigned an “A”, “B”, or “C” designation, with the clade grouping with the reference sequence being given the designation “A”, and the other clade designated “B”. If there were three clades in total, then the non-reference clade with the fewest sequences was designated “C” (**Figure 3, Supplementary Figure 1**).

**Figure 2.**
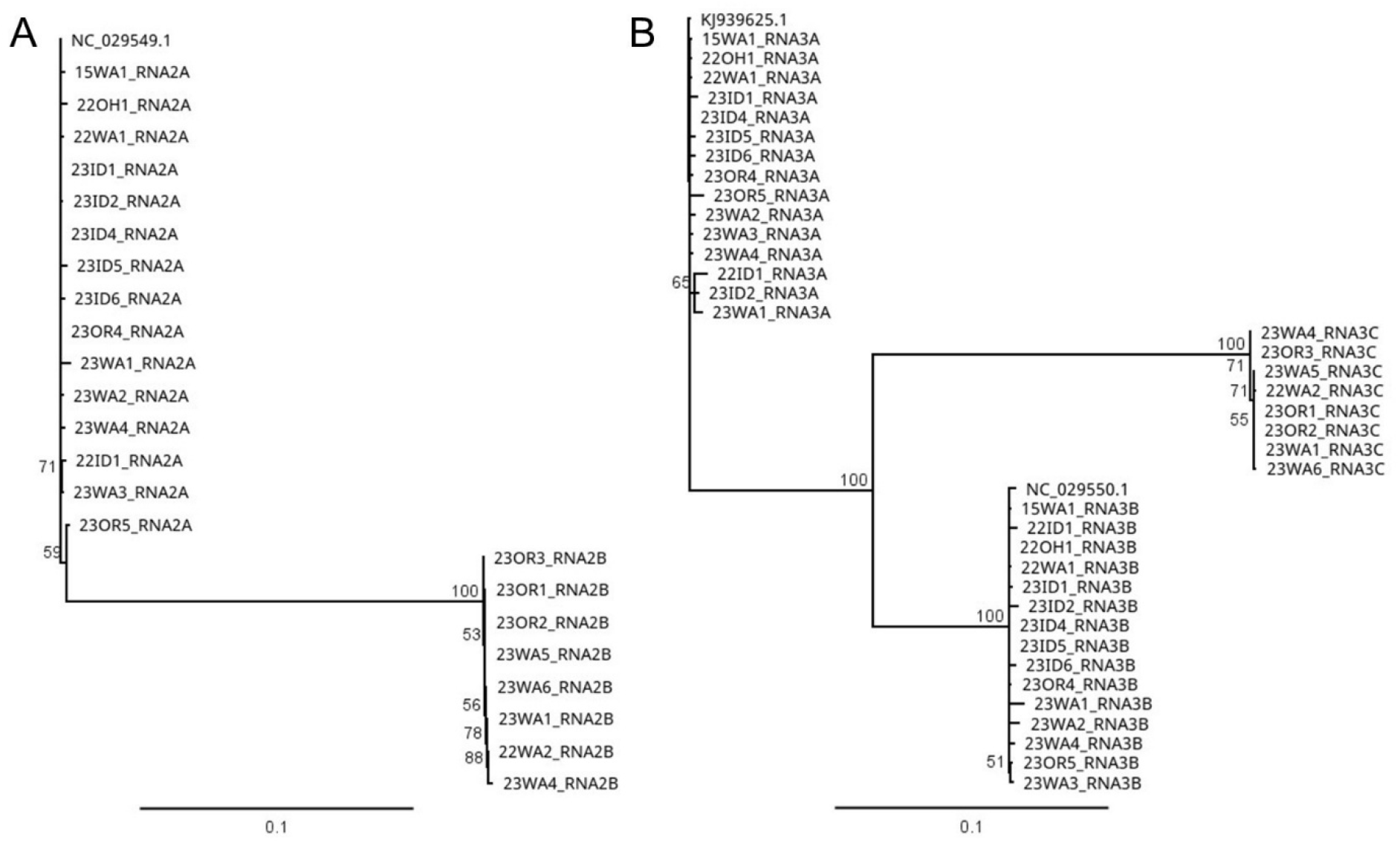
Phylogenetic analysis reveals two or three main clades for most RNA segments of HPWMoV isolates. Shown as an example are RNA2 with two clades (**A**) and RNA3 with three clades (**B**). The clade containing the reference sequence is given the designation “A”, as in “RNA2A”, and the non-reference clade is designated “B” (i.e., “RNA2B”). RNA3 was already known to possess A and B variants, and the clades are named accordingly. The third clade, which does not contain the RNA3A or RNA3B reference sequences, has been designated RNA3C. Both panels are roughly to the same horizontal scale. Shown at the nodes are bootstrap values out of 100 replicates. Phylogenetic trees for the remaining RNA segments can be found in Figure 3 (RNA6 and RNA8) and **Supplementary** Figure 1 (all others).

**Figure 3.**
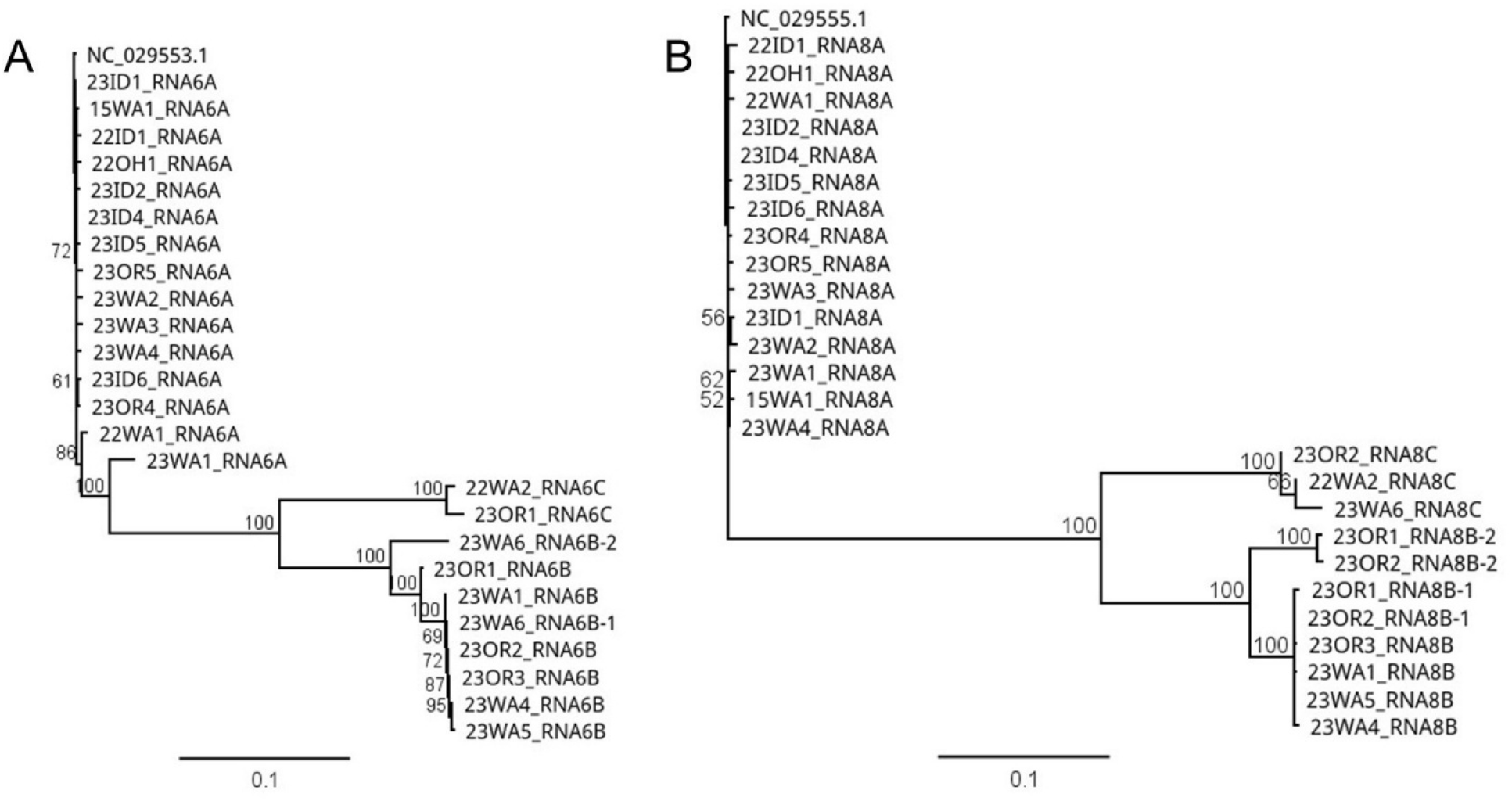
Some RNA segments show further sequence divergence within a clade of HPWMoV isolates. Shown as an example are RNA6 (**A**) and RNA8 (**B**). Sequences with a “-1” or “-2” suffix indicate two subvariants of the same RNA segment (i.e., RNA6B or RNA8B) from the same sweet corn sample. The clade containing the reference sequence is given the designation “A” as in “RNA6A” and the non-reference clades are designated “B” and “C” (i.e. “RNA6B”), with the non-reference clade with the fewest sequences receiving the “C” designation. Both panels are roughly to the same horizontal scale. Shown at the nodes are bootstrap values out of 100 replicates. Phylogenetic trees for the remaining RNA segments can be found in Figure 2 (RNA2 and RNA3) and **Supplementary** Figure 1 (all others).

Multiple sequence alignment and pairwise nucleotide identity values between these RNA segment variants supported these phylogenetic groupings (**Figure 4, Supplementary Table 2**), with clear differences in genetic distance between clades versus within clades. For instance, all RNA2A sequences shared on average 99.72% sequence identity, and all RNA2B sequences were 99.92% identical to one another, but pairwise comparison of RNA2A and RNA2B sequences showed only 85.67% nucleotide identity, on average. From here on, the different versions of the same RNA segment are called “variants,” i.e., RNA2A and RNA2B are two variants of the RNA2 genome segment.

**Figure 4.**
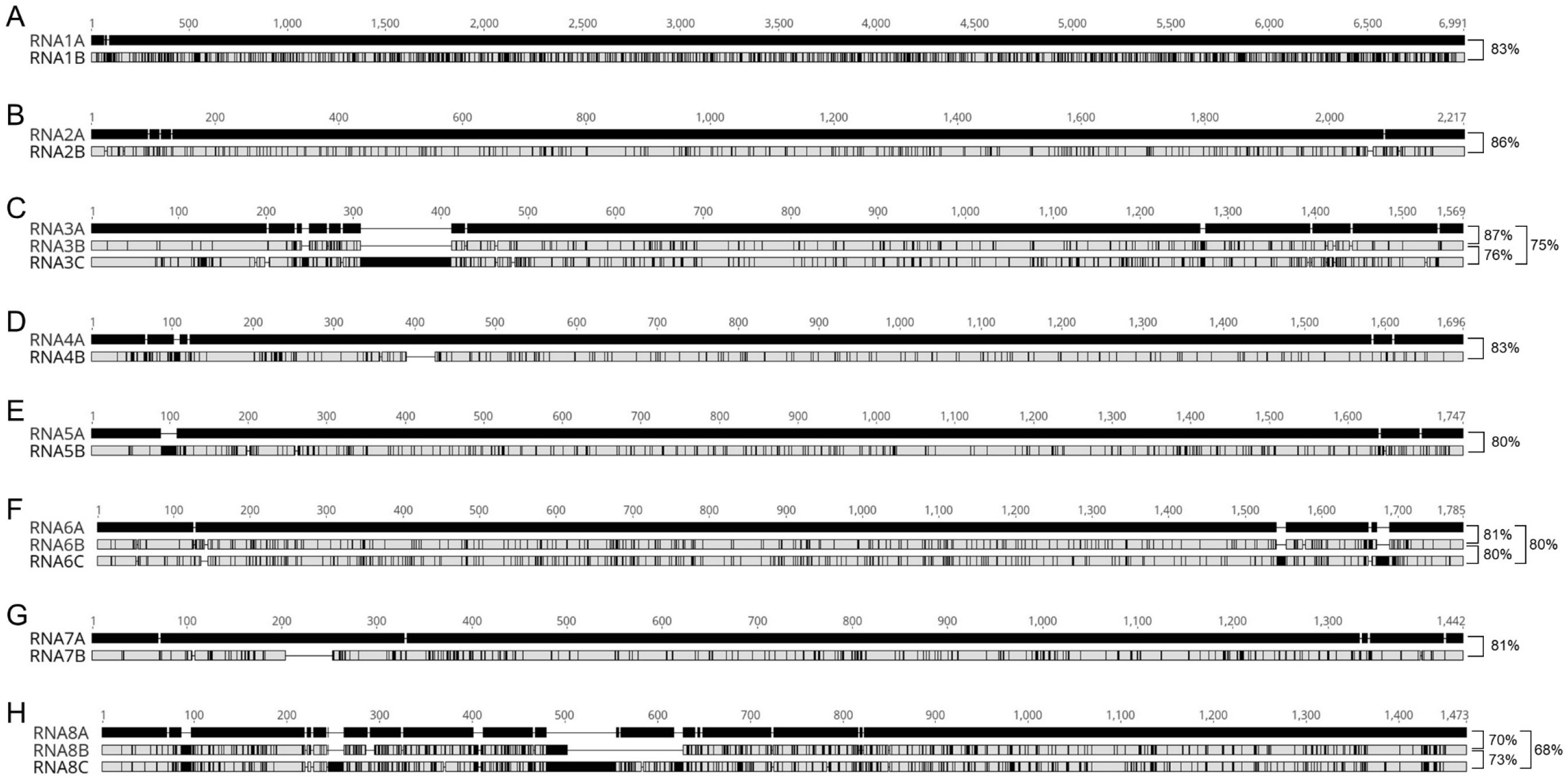
There are two or three major variants for each RNA segment of HPWMoV. Shown is an alignment of consensus sequences of the “A,” “B,” and “C” variants for each RNA segment. Consensus sequences for each variant (e.g., RNA1A) were generated by aligning all sequences of that variant that share >90% sequence identity. Then, the two (A & B) or three (A, B, & C) consensus sequences for each RNA segment were aligned, with the “A” variant designated as the reference sequence (black). Differences from the reference are indicated with vertical black lines. The percent nucleotide identity between variants is indicated at the end of the sequences with brackets

Phylogenetic grouping and nucleotide identity thresholds were not clear cut for all RNA segments. For some segments, there was further sequence divergence within the clades, e.g., for RNA6B and RNA8B (**Figure 3**). In the case of RNA6B, two different subvariants of RNA6B were present within the 23WA6 sample. One subvariant closely resembled the other RNA6B sequences, with an average pairwise nucleotide identity of 99.53% (**Supplementary Table 2**). However, the other subvariant, designated 23WA6_RNA6B-2, only shared an average of 93.5% nucleotide identity with the other RNA6B sequences. However, this divergence did not approach the genetic distance observed between RNA6B and RNA6A (81.27%) nor the average pairwise nucleotide identity between RNA6B and RNA6C (80.29%), although this particular RNA6B subvariant was slightly more similar to RNA6C than the other RNA6B sequences with 82.84% nucleotide identity, on average. Similarly, two subvariants of RNA8B were present in samples 23OR1 and 23OR2. Each sample contained a typical RNA8B subvariant (23OR1_RNA8B-1, 23OR2_RNA8B-1) with 99.66% identity with the other RNA8B sequences, and a second subvariant (given the suffix “-2”, **Figure 3**) that shared only 92.92% nucleotide identity, on average, with other RNA8B sequences. Interestingly, these two divergent RNA8B subvariants were 99.32% similar to one another. As observed for RNA6B, these divergent subvariants did not approach the level of sequence divergence present between RNA8A and RNA8B (69.99%) or RNA8B and RNA8C (67.64% pairwise nucleotide identity, on average). These two samples were collected from adjacent fields in Morrow County, Oregon, about three-quarters of a mile from a dryland wheat cover crop. Both samples were also co-infected with WSMV (**Table 1**).

Multiple sequence alignment showed that, in addition to many nucleotide differences between variants of the same RNA segment, there were also significant insertions and deletions. The B variants of RNA1 and RNA2 differed from the A variants by only a few small indels and were roughly the same length. However, RNA3C had a notable 100 bp insertion in the first third of the sequence relative to RNA3A and RNA3B (**Figure 4**). The indels between variants of the other RNA segments were smaller, with a 35 bp deletion in the first part of RNA4B relative to RNA4A, a 20 bp insertion in the beginning of RNA5B, two small 12-16 bp insertions towards the end of RNA6C, and a 49 bp in the first part of RNA7B. RNA8 has many more indels than the other RNA segments. RNA8B and RNA8C shared a 10 bp insertion towards the beginning of the sequence. Further down the sequence, RNA8B was missing a 63 bp stretch present in the other two variants, but contained a 23 bp insertion relative to RNA8A in the same region shared with RNA8C. However, the RNA8C insertion was more extensive, and extended 52 bp beyond the piece shared with RNA8B. Interestingly, the alternative RNA8B subvariants 23OR1_RNA8B-2 and 23OR1_RNA8B-2 both shared a 16 bp insertion with RNA8C that was not observed in the other RNA8B sequences. Multiple sequence alignments of all sequences of each genome segment can be found in **Supplementary Figures 2-5**, with nucleotide-level alignments in FASTA format provided as supplementary files.

### 3.4 HPWMoV isolates sort into two major types with similar RNA segment composition

While two to three variants were found for each RNA segment, for a total of 19 possible unique segments, each sample contained just 10 genome segments, on average (nine excluding the outliers 23WA1 and 23WA4, **Supplementary Table 3**). Considering HPWMoV is an octopartite virus, in general, each sample contained at least one variant of all RNA segments with one to two additional segment variants. However, there were distinct differences across RNA segments are far as how many variants were observed in a single sample. The most common RNA segment to have multiple variants within a sample was RNA3, with 71% of samples containing more than one RNA3 variant, which was always a combination of RNA3A and RNA3B when two variants of RNA3 were present. In contrast, the RNA segments least likely to be present as more than one variant were RNA1, RNA2, RNA4, and RNA7; except for the co-infected samples 23WA1 and 23WA4, no other samples in our study had more than one variant of RNA1, RNA2, RNA4, or RNA7. Besides the co-infected samples, more than one variant of RNA5 was observed in four samples (15WA1, 22OH1, 23OR1, 23OR4). However, as many as three variants of RNA8 were found in a single sample in our study and, in one instance, RNA8C seemed to replace RNA8B as it was the only variant of RNA8 found in sample 23WA6. Similarly, RNA6C seemed to replace RNA6B in isolate 22WA2.

The distribution of RNA segment variants was not random, and within each sample they generally sorted into two major types, with a few minor exceptions (**Figure 5**). The first type, which we have designed type A, contains the A variant of all eight genome segments, plus RNA3B. This was the predominant variant configuration found in the samples, with 13 samples showing this exact pattern of RNA segments. Three samples presented this pattern with the addition of RNA5B (15WA1, 22OH1, 23OR4, **Supplementary Figure 5A**). These three isolates were also designated as type A, for a total of 16 type A isolates in this study.

**Figure 5.**
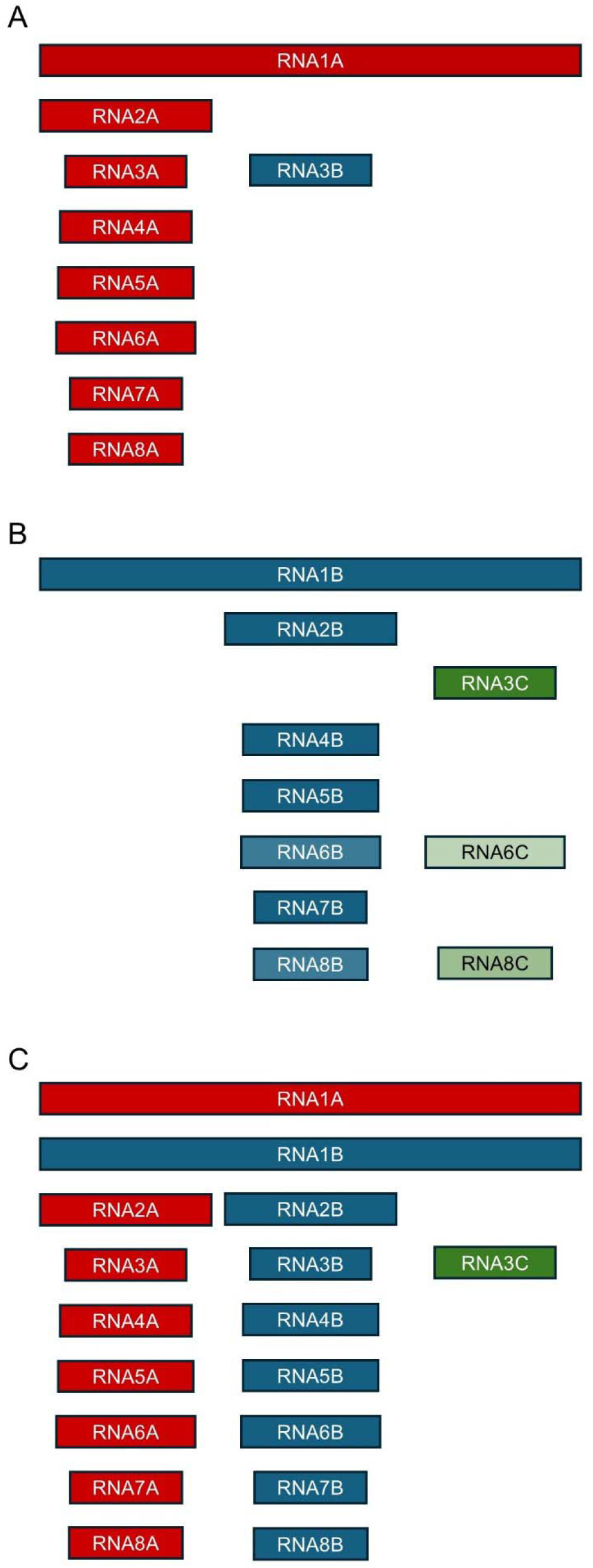
HPWMoV isolates sorted into two major types with potential instances of co-infection. Shown is a visual representation of type A (A), type B (B), and type A+B (C) isolates. Genome segments are represented by rectangles, which are to scale. “A” variants are red, “B” variants are blue, and “C” variants are green. The transparency of the coloring of each segment represents its relative appearance in that isolate type. For instance, RNA3B is always present in type A isolates whereas RNA6C was only present in 2 of 6 type B isolates in the current study. Type A+B was only found in the potentially co-infected samples 23WA1 and 23WA4. Both samples had the same genome segment composition. A detailed account of the abundance of each RNA segment variant in every sample can be found in Supplementary Table 3. A visual representation of minor exceptions on type A and type B isolate composition observed in the study can be found in **Supplementary** Figure 5.

The other major variant configuration observed, which we designated as type “B”, generally included the B variant for all RNA segments except RNA3, which was presented as the RNA3C variant (**Figure 5B**). Three of the samples contained this exact configuration. In sample 23WA2 that otherwise followed the type B pattern, the RNA6B variant was not present and, instead, the only form of RNA6 observed was RNA6C. Similarly, sample 23WA6 generally followed the type B configuration, but RNA8 was only present as RNA8C. One other minor exception was sample 23OR1 which followed the typical type B pattern but additionally had RNA5A (**Supplementary Figure 5B**). All of these isolates were designated type B, for a total of six type B isolates. Interestingly, the C variants of RNA6 and RNA8 were only observed in type B isolates (**Supplementary Table 3**).

Curiously, samples 23WA1 and 23WA4 contained the variant composition of both type A and type B isolates, for a total of 17 genome segments (**Figure 5C**). In fact, they contained every RNA segment variant found in this study except for RNA6C and RNA8C. The presence of two variants for RNA1, RNA2, RNA4, and RNA7 and all three variants for RNA3 was confirmed in both samples via Sanger sequencing (**Supplementary Table 1**). These two isolates have been designated type A+B. It is unknown how many plants were sampled for isolate 23WA4 so this could be the result of sampling a plant infected with a type A isolate and another with a type B isolate in the same field. However, sample 23WA1 was collected from a single plant and may represent co-infection of a plant by two HPWMoV isolates of differing types.

Both type A and type B isolates were collected from the Columbia Basin in Washington and Oregon, as well as both combined type A+B samples. In contrast, only type A isolates were collected from the Treasure Valley in Idaho and Oregon (**Figure 1**).

### 3.5 All but one of the previously sequenced HPWMoV isolates aligned with the two major isolate types

To determine how previously sequenced HPWMoV samples align with the two isolate types designated in this study, we performed multiple sequence alignments between all previously sequenced HPWMoV isolates and the consensus sequences for each RNA variant from our 21 isolates (**Table 3, Supplementary Table 4**). The type isolate (Nebraska) was excluded from the analysis as it defines type A. There were 17 previously sequenced samples in total, but many were incomplete and three were confirmed composite samples comprised of multiple plants (CoPhil, HPWMoV_NWB1, and HPWMoV_NWB2). An alignment was performed for each RNA segment separately (see **Supplementary Files**) and an “A,” “B,” or “C,” designation was given based on percent nucleotide identity to the consensus sequences. In most cases, >90% nucleotide similarity was found between a previously sequenced RNA segment and one of our consensus sequences. Then, the variant assignments of each RNA segment were taken collectively to classify the isolate as type A or type B. Five of the previously sequenced isolates (GG1, HP2W, HPWMoV_ID, HPWMoV_MI, and KS7) presented the typical type A configuration of all A variants plus RNA3B (**Figure 5A**). Five other isolates (BCHPV1, BCHPV2, HP1G, HPWMoV_ID, and HPWMoV_MI) also received the type A designation with the only deviation from the typical type A pattern being the presence of RNA5B rather than RNA5A. Isolate K1 presented the typical type B pattern (all B variants except for RNA3C, **Figure 5B**). Four other isolates (H1, HPWMoV_NWB2, TT2025-3, and W1) had an alternate type B isolate configuration with RNA8C rather than RNA8B, and two of these isolates had RNA6C instead of RNA6B, similar to what has been observed for some type B isolates in this study. Isolate TT2025-31 had the same configuration but with a second RNA5B and two additional RNA6 segments. The isolate BCWS5 could not be assigned to a type due to so few RNA segments being sequenced, but the sequenced segments were designated RNA3A, RNA3B, and RNA6A, indicating a likely type A isolate.

**Table 3.**
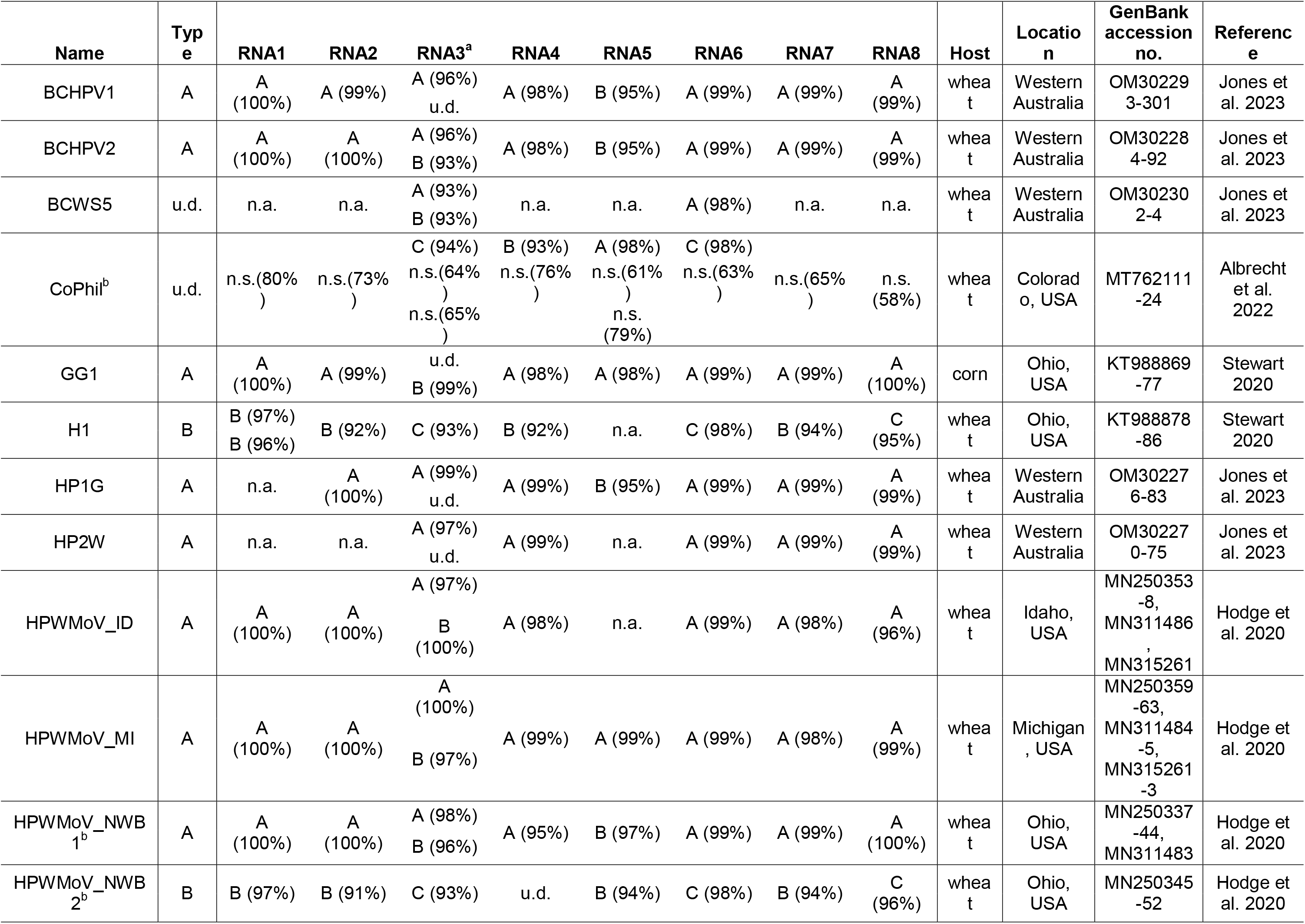
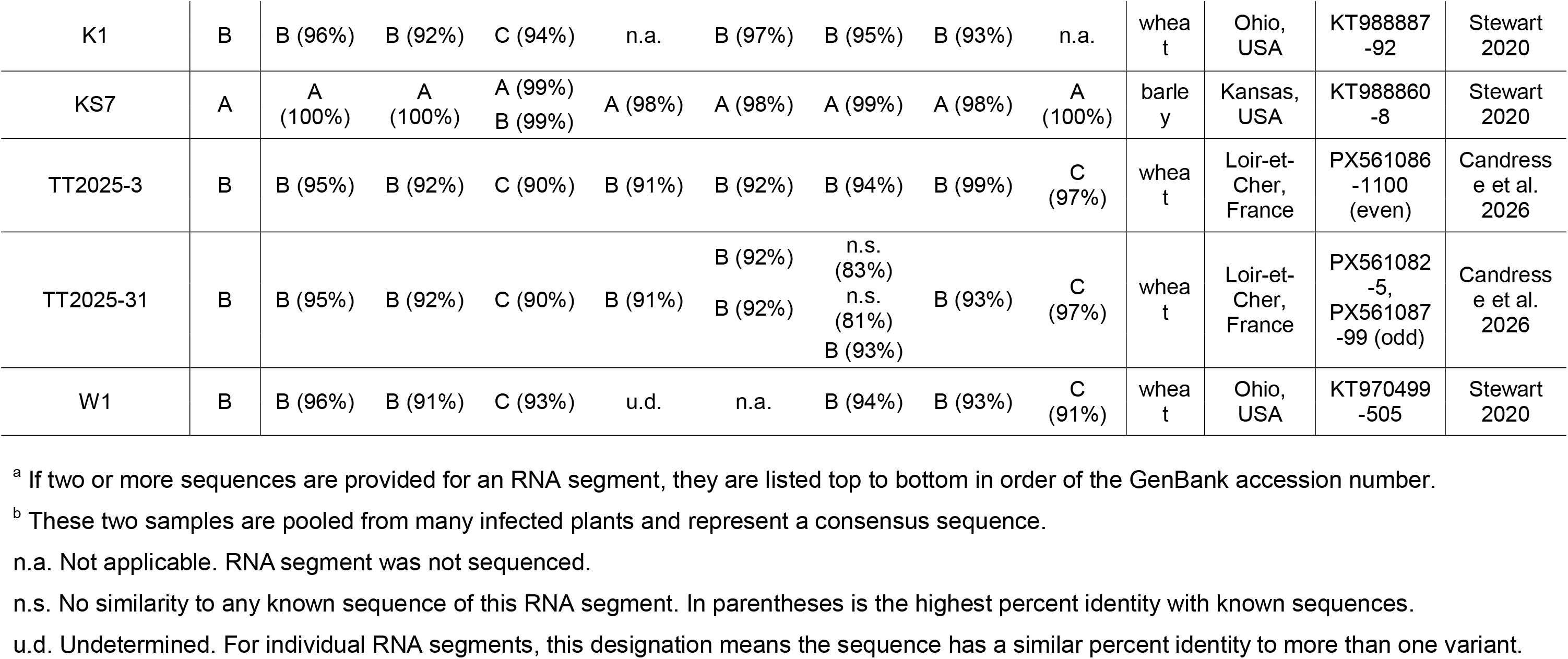
Isolate type and RNA variant assignment of previously sequenced HPWMoV samples based on percent nucleotide identity to the RNA segment consensus sequences generated from 21 HPWMoV isolated sequenced in this study.

The number of variants for each genome segment in previously sequenced isolates also aligned with what was observed in this study. While RNA3 was often present as two variants, only one variant was observed for RNA2, 4, 7, and 8, with the exception of composite sample CoPhil. Two variants of RNA1 were detected in sample H1, and two variants of RNA5 and three variants of RNA6 were all detected in sample TT2025-31. Interestingly, both RNA5 variants from TT2025-31 were assigned to RNA5B, making this isolate similar to those in our study that had two subvariants of RNA6B and RNA8B. Of the three RNA6 variants sequenced from TT2025-31, two could not be assigned to any known variant of RNA6 (A, B, or C) and the two sequences differed from one another, sharing only 80% nucleotide identity. These sequences may belong to two new clades of RNA6. Besides these, multiple variants of other RNA segments may have also been present in samples from previous studies but just not sequenced.

The only isolate that did not neatly align with the two isolate types is CoPhil, which was a composite sample collected from wheat plants in Phillips County, Colorado (Albrecht et al., 2022). Of the 14 genome segments submitted to GenBank from the CoPhil sample, only four could be assigned to known variants (RNA3C, RNA4B, RNA5A, and RNA6C), with no close nucleotide similarity for any of the other segments which only had 58-79% nucleotide similarity to any other HPWMoV sequences. Partial sequences for RNAs 1, 2, 5, 7, and 8 were also obtained from the sample but not deposited to GenBank. However, based on the closest BLAST hit for each of these undeposited sequences, as reported by the authors, they most likely corresponded to RNA1B (closest match is RNA1 from type B isolate HPWMoV_NWB2), RNA2B (closest to RNA2 of type B isolate W1), RNA5B (closest to RNA5 from a pool of type B isolates H1, K1, W1), RNA7B (closest to RNA7 of HPWMoV_NWB2) and RNA8C (closest to RNA8 of isolate H1). This collection of RNA segments indicated that sample CoPhil is likely a type B isolate with potential co-infection with a hitherto yet underdetermined isolate type or emaravirus species, if the deposited sequences are not misassembled. Indeed, the deposited sequences that could not be assigned to a HPWMoV genome segment had an average reported coverage of 14.8 reads, with more than half having fewer than 10 reads. The deposited sequences also lacked the conserved sequences at the end of each genome segment that are shared across emaravirus species. In some alignments, the CoPhil sequences extended beyond where the conserved ends would be, though the same is true of many of the previously sequenced isolates (see **Supplementary Files**).

There was no discernable pattern of isolate type by plant host. Both A and B type isolates were previously sequenced from wheat, and both types were sequenced from sweet corn in this study. The only sample sequenced from barley to date is a type A isolate. All isolates sequenced from Western Australia thus far are type A isolates, and both isolates sequenced from France are type B. Both isolate types were collected from various regions of the United States (**Table 3**).

### 3.6 Only three recombination events were predicted among the 21 isolates sequenced in this study, two in RNA5 and one in RNA7

To detect potential recombination events, the alignments of all the sequences generated in this study were analyzed with RDP5. Three recombination events were predicted: two in RNA5 and one in RNA7 (**Table 4**). Predicted recombination events are only reported if they were detected by seven out of the nine detection algorithms. Since chimeric assembly of short reads can lead to artificial recombination signatures, to rule out such misassemblies, primers were designed flanking each recombination event and the resulting PCR products sequenced via Oxford Nanopore (ONP) long-read sequencing. Two partially overlapping sequences were obtained for RNA5B, covering the two predicted recombination events.

**Table 4.**
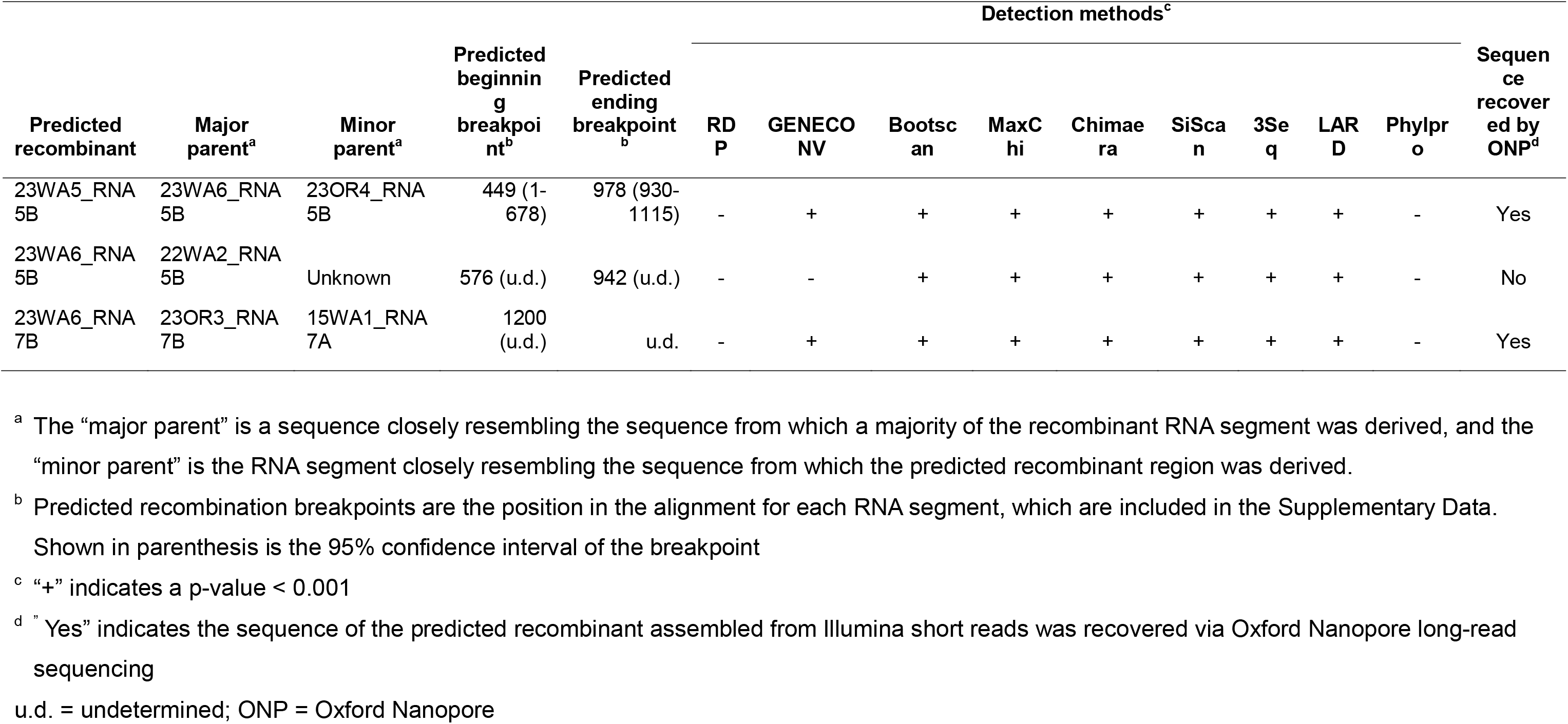
Predicted recombination events among HPWMoV sequences generated in this study.

The *de novo* assembled sequence for 23WA5_RNA5B was confirmed, with the two long-read sequences matching the Illumina assembled sequence almost exactly, except for a single base which was considered “low confidence” in the ONP sequencing, where the alternate base was detected at a slightly lower frequency than the base called (see **Supplementary Data**). On the contrary, the assembled sequence for 23WA6_RNA5B was not detected, as the two sequences obtained by ONP only shared 98% similarity with the Illumina assembly and only 99% similarity with one another. It is unclear whether the differences between the two ONP sequences are due to sequencing errors or the presence of multiple unique RNA5B sequences. Indeed, it is possible that there were additional versions of RNA5B in this sample, and the one assembled from Illumina short reads was different from the one(s) amplified prior to ONP sequencing. Alternatively, the *de novo* assembled sequence is a chimeric assembly of two versions of RNA5B. Either way, misassembly could not be ruled out as a potential cause of the recombination signature in the 23WA6_RNA5B sequence. Considering that 23WA6_RNA5B was identified as the major parent for the predicted recombination event in 23WA5_RNA5B, it is likely that the chimeric region of 23WA6_RNA5B (whether due to recombination or misassembly) is also the reason for this predicted recombination event, and the recombinant sequence was misidentified as the major parent, as is common in RDP5 analyses (Martin, 2020). The similarity in predicted breakpoints between the two predicted recombination events in RNA5B corroborates that this may indeed be a single event.

As for the predicted recombination event in RNA7, the sequence of 23WA6_RNA7B was confirmed by long-read sequencing, as the sequence derived from ONP matched the Illumina assembly perfectly, except for the first nucleotide. Visual examination of the alignment did not reveal any differences between 23WA6_RNA7B and the RNA7B consensus sequence in the predicted recombination region or in the entire RNA7 sequence. Of the predicted parents, 15WA_RNA7A had the most differences from the consensus sequence, and RDP5 gave a warning that 15WA1_RNA7B may be the actual recombinant. The 15WA1_RNA7A sequence was also confirmed by ONP sequencing, ruling out chimeric assembly as the cause of the recombination signature.

### 3.7 Newly designed HPWMoV primer and probe sets identified all 20 isolates tested in diagnostic assays and likely all HPWMoV samples sequenced to date based on *in silico* analysis

Two new primer and probe sets for RT-qPCR were designed to improve seed health testing diagnostics, one targeting RNA2 and the other targeting RNA4. The RNA2 primer and probe set was designed prior to sample collection from the Columbia Basin and Treasure Valley, and was based on the RNA2 sequences of previously sequenced HPWMoV isolates. The RNA4 primer and probe set was designed based on the sequences of the 21 new HPWMoV isolates sequenced in this study. Both *in silico* prediction and actual RT-qPCR tests in duplex were performed for both primer and probe sets with 20 of the 21 isolates from this study. Only isolate 15WA1 was excluded from RT-qPCR tests due to a lack of plant tissue. Leaf tissue for all isolates was tested as well as seed tissue for four isolates. Only *in silico* analysis could be performed on previously sequenced isolates.

For RNA2, *in silico* analysis of the 21 isolates sequenced in this study (**Figure 6A**) only showed a single mismatch in each of the primers for some type B isolates. There were no mismatches in the probe annealing site for any of the RNA2A sequences, and only one mismatch in the middle of the forward primer for isolates containing RNA2B and one mismatch in the reverse primer for RNA2B sequences, plus a mismatch for isolate 23WA2 in the same position. Since a single mismatch can easily be tolerated in a primer, these single mismatches were not predicted to lead to missed detection of these isolates. This prediction was confirmed via qRT-PCR testing of the RNA2 primer and probe set in duplex, with all 20 isolates recognized (**Table 5**). The average Cq value was lower for leaf tissue overall than seed tissue (18.14 ± 2.35 standard deviation vs. 25.05 ± 1.27). The RNA2 primer and probe set seemed to have a slight bias for type A isolates (average Cq of 16.92 ± 1.35 for type A vs. 21.11 ± 1.40), which may be due the single mismatch in the reverse primer is only present in type B isolates, but further testing with greater sample sizes would be needed for confirmation. The RNA2 primer and probe set gave no amplification from non-infested seed and no template control samples but sometimes had very late non-specific amplification in non-infected leaf tissue (**Table 5**). For this reason, a Cq cut-off value of 35 is proposed but final cut-off values to be used for seed health testing can be determined through ring testing and further validation of this assay.

**Figure 6.**
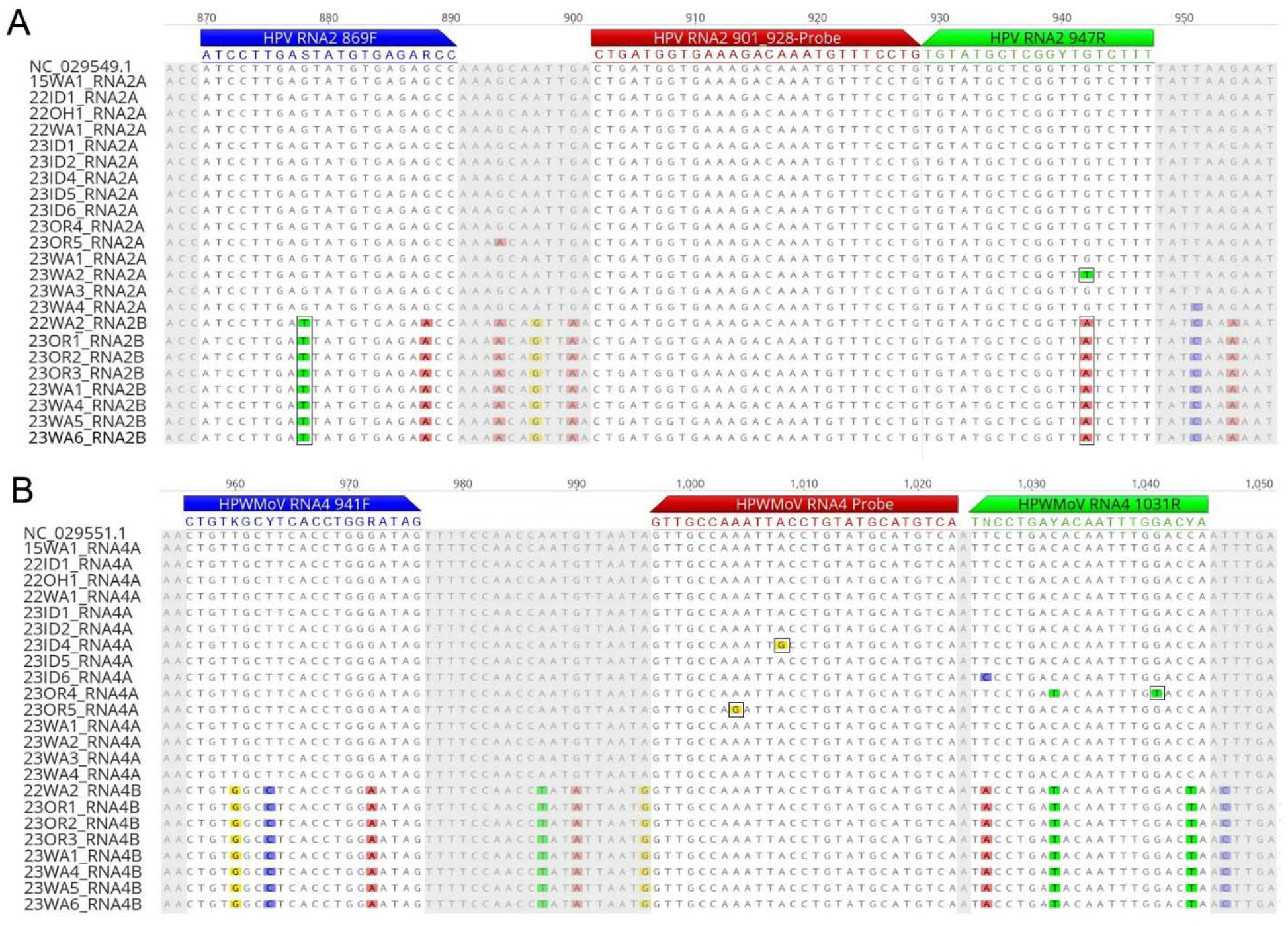
Both the RNA2 and RNA4 primer and probe sets are predicted *in silico* to recognize all 21 HPWMoV isolates sequenced in this study. Pictured are the multiple sequence alignments for RNA2 and RNA4 in the region where the primer and probe sets anneal. Forward primers are shown in blue, reverse primers in green, and probes in red. Nucleotide differences from the consensus are highlighted. Differences that do not match the primer or probe sequences are outlined with a black box. Nucleotides not in the region where primers or probes anneal have been grayed out. The sequences for the reverse primer and the RNA4 probe are reverse complements so that mismatches could be identified more easily as only the forward strand of DNA is presented here.

**Table 5.**
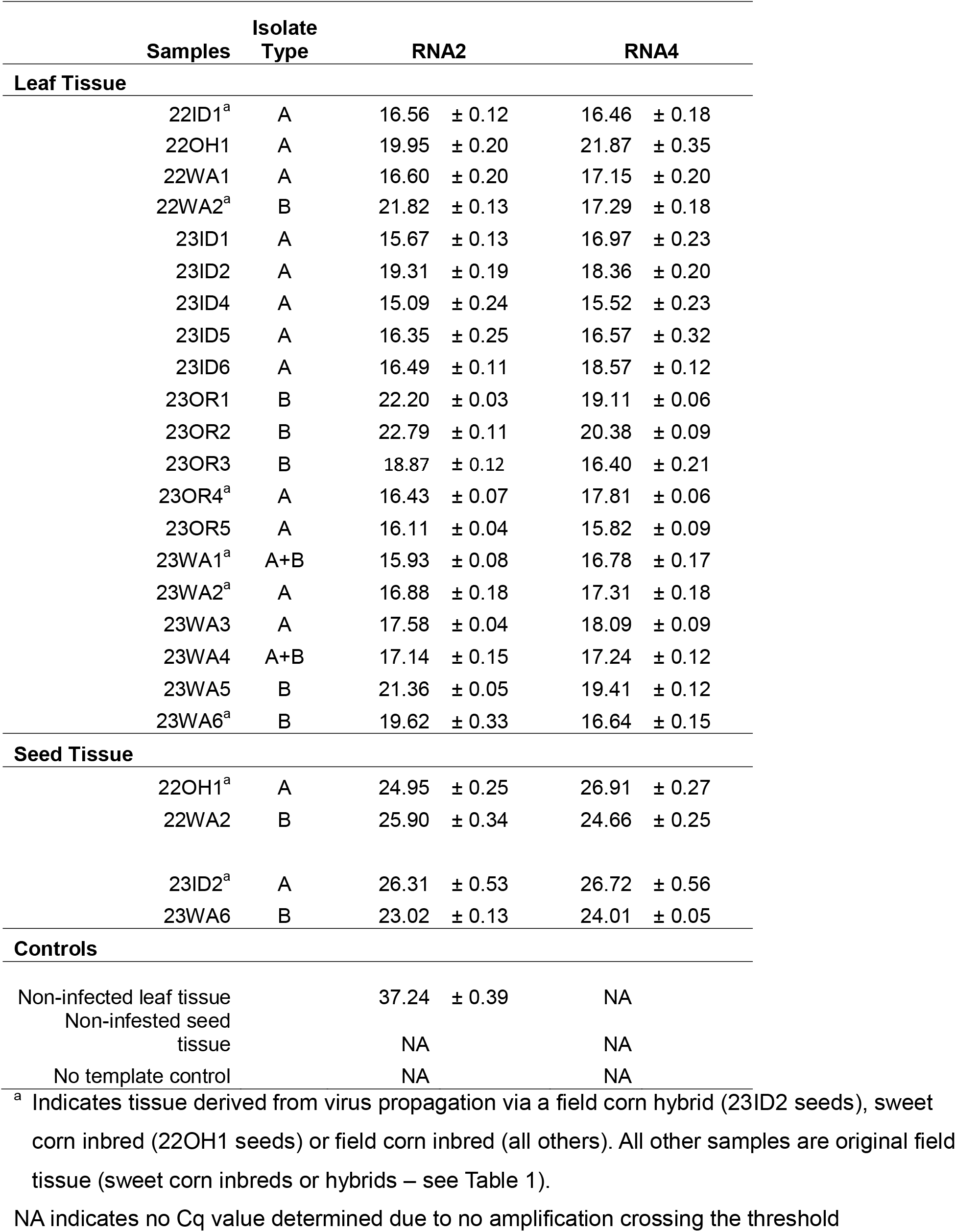
Summary of Cq values and standard deviation among three technical replicates obtained from RT-qPCR tests of 20 isolates of HPWMoV sequenced in this study.

| Samples | Isolate Type | RNA2 |  | RNA4 |  |
| --- | --- | --- | --- | --- | --- |
| Leaf Tissue |  |  |  |  |  |
| 22ID1<sup>a</sup> | A | 16.56 | ± 0.12 | 16.46 | ± 0.18 |
| 22OH1 | A | 19.95 | ± 0.20 | 21.87 | ± 0.35 |
| 22WA1 | A | 16.60 | ± 0.20 | 17.15 | ± 0.20 |
| 22WA2<sup>a</sup> | B | 21.82 | ± 0.13 | 17.29 | ± 0.18 |
| 23ID1 | A | 15.67 | ± 0.13 | 16.97 | ± 0.23 |
| 23ID2 | A | 19.31 | ± 0.19 | 18.36 | ± 0.20 |
| 23ID4 | A | 15.09 | ± 0.24 | 15.52 | ± 0.23 |
| 23ID5 | A | 16.35 | ± 0.25 | 16.57 | ± 0.32 |
| 23ID6 | A | 16.49 | ± 0.11 | 18.57 | ± 0.12 |
| 23OR1 | B | 22.20 | ± 0.03 | 19.11 | ± 0.06 |
| 23OR2 | B | 22.79 | ± 0.11 | 20.38 | ± 0.09 |
| 23OR3 | B | 18.87 | ± 0.12 | 16.40 | ± 0.21 |
| 23OR4<sup>a</sup> | A | 16.43 | ± 0.07 | 17.81 | ± 0.06 |
| 23OR5 | A | 16.11 | ± 0.04 | 15.82 | ± 0.09 |
| 23WA1<sup>a</sup> | A+B | 15.93 | ± 0.08 | 16.78 | ± 0.17 |
| 23WA2<sup>a</sup> | A | 16.88 | ± 0.18 | 17.31 | ± 0.18 |
| 23WA3 | A | 17.58 | ± 0.04 | 18.09 | ± 0.09 |
| 23WA4 | A+B | 17.14 | ± 0.15 | 17.24 | ± 0.12 |
| 23WA5 | B | 21.36 | ± 0.05 | 19.41 | ± 0.12 |
| 23WA6<sup>a</sup> | B | 19.62 | ± 0.33 | 16.64 | ± 0.15 |
| Seed Tissue |  |  |  |  |  |
| 22OH1<sup>a</sup> | A | 24.95 | ± 0.25 | 26.91 | ± 0.27 |
| 22WA2 | B | 25.90 | ± 0.34 | 24.66 | ± 0.25 |
| 23ID2<sup>a</sup> | A | 26.31 | ± 0.53 | 26.72 | ± 0.56 |
| 23WA6 | B | 23.02 | ± 0.13 | 24.01 | ± 0.05 |
| Controls |  |  |  |  |  |
| Non-infected leaf tissue |  | 37.24 | ± 0.39 | NA |  |
| Non-infested seed tissue |  | NA |  | NA |  |
| No template control |  | NA |  | NA |  |

<sup>a</sup> Indicates tissue derived from virus propagation via a field corn hybrid (23ID2 seeds), sweet corn inbred (22OH1 seeds) or field corn inbred (all others). All other samples are original field tissue (sweet corn inbreds or hybrids – see Table 1).
NA indicates no Cq value determined due to no amplification crossing the threshold

As for predicted recognition of previously sequenced isolates by the RNA2 primer and probe set (**Supplementary Figure 6A**), there was a single mismatch halfway through the reverse primers for isolates H1 and W1, and a single mismatch towards the 3’ end of the probe among these two isolates as well as HPWMoV_NWB2, K1, and T2025-3. There was a single mismatch at the 5’ nucleotide of the probe for isolate TT2025-31. Mismatches are not well tolerated in the probe and could lead to a missed detection, but this was not the case for the RNA4 probe and isolates 23ID4 and 23OR5, which each contained a single mismatch in the 3’ half of the probe (**Figure 6B**) but were still detected in qRT-PCR tests (**Table 5**). Of these isolates, all but TT2025-31 were predicted to be recognized by the RNA4 set and so would still test positive for HPWMoV in the duplex assay. The one isolate with many mismatches that would likely lead to a missed detection was RNA2 from the CoPhil isolate (MT762124), but this sequence may belong to another emaravirus species due to its very low similarity to other HPWMoV RNA2 sequences, or could be a misassembly. Furthermore, one of the RNA4 segments of the CoPhil isolate was predicted to be recognized by the RNA4 primer and probe set, so the CoPhil sample would be HPWMoV positive in the duplex assay.

For RNA4, *in silico* analysis of the 21 isolates from this study (**Figure 6B**) indicated only a few potential mismatches within the primers and probe. There was a single mismatch to isolate 23OR4 in the 5’ region of the reverse primer, but considering its position, this likely would not lead to a missed detection, which was confirmed in the RT-qPCR tests. There was a single mismatch in the probe for each of 23ID4 and 23OR5 but actual tests of the RNA4 primer and probe set on these isolates were effective, producing a low Cq value of 15.52 ± 0.23 and 15.82 ± 0.09, respectively. Indeed, all 20 isolates tested were recognized in the RT-qPCR tests (**Table 5**) with an average Cq of 17.69 ± 1.53 for leaf tissue and an average Cq of 25.57 ± 1.26 for seed tissue. The RNA4 primer and probe set seemed to have similar detection of the two isolates types, with an average Cq close to 18 for both isolate types. Amplification was not detected from the non-infected leaf, non-infested seed, and no template negative controls.

For previously sequenced HPWMoV isolates (**Supplementary Figure 6B**), there was a single mismatch towards the 5’ end of the probe for isolate TT2025-31 which, based on its position, could cause a missed detection. RNA4A of the CoPhil sample (MT762119) had 5 mismatches in the RNA4 probe, 2 mismatches in the forward primer and 3 mismatches in the reverse primer, which would certainly lead to a missed detection. However, the CoPhil sample also had an RNA4B sequence which perfectly matched the RNA4 primer and probe set. Hence, if the CoPhil sample was tested in the RNA4 qRT-PCR assay, the sample would likely be considered positive for HPWMoV. Among other previously sequenced isolates, there was one mismatch in the 5’ half of the forward primer for isolate HPWMoV_NWB2, as well as a single mismatch in the 5’ region of the reverse primer for isolates BCHPV2 and HP1G. These single mismatches, especially in the 5’ portion of the primers, were not predicted to lead to missed detections. No prediction could be made for detection of isolate HPWMoV_NWB1 with the RNA4 primer and probe set as there was no sequence information available for this region of RNA4, though this isolate was predicted to be recognized by the RNA2 set with no mismatches.

## 4. Discussion

This study presents the most complete picture of HPWMoV sequence diversity to date and proposes a classification scheme for HPWMoV isolates that captures all but one previously sequenced isolate. The presence of two major isolate types was first observed by Stewart (2016) who referred to them as “Nebraska-like” or “Ohio-like” depending on whether they most closely matched the reference sequence from Nebraska or the wheat isolate sequences from Ohio, which correspond to type A and type B, respectively, as defined in this study. Previous studies have also reported that HPWMoV isolates phylogenetically form two groups, though in those studies the reference clade was termed “Group B” (Jones et al., 2023; Pozhylov et al., 2022). We maintain that the reference clade and isolate type should be designated “A,” which we now know represents the most common HPWMoV isolate type around the world among samples sequenced to date. Importantly, our isolate type designation is based on a shared complement of genome segments, not phylogenetic grouping of individual genome segments. The genome segments themselves may group into two or three clades with some further divergence, yet they still sort into only two isolate types. Hence, the isolate type designation is broader and more encompassing than examining any single RNA segment. While previously the two major types of HPWMoV were referred to as strains, further characterization is needed to determine if they have sufficiently distinct biological properties to designate them as unique strains. The two isolate types are not serologically distinct, as both types are recognized by the same commercial enzyme-linked immunosorbent assays.

The deep sequencing in this study allowed us to observe multiple variants for every genome segment which had only been observed once before for HPWMoV, in the composite sample CoPhil, which was sequenced from a pool of plants (Albrecht et al., 2022). Here, we report two variants of RNAs 2, 4, and 7, and three variants of RNA8 in a single plant for the first time, as well as the first report of two complete HPWMoV genomes from a single plant. The “co-infections” we observed have also been observed for the pentapartite emaravirus palo verde broom virus (Adegbola et al., 2025), showing there is precedence for two emaravirus isolates of the same species infecting the same plant. Our approach of performing enrichment for virus-like particles prior to sequencing all samples also differed from most previous studies and allowed us to obtain a high sequencing depth while also ensuring the vast majority of the viral sequences were packaged into virions and were not defective or aberrant variations of the viral genome that are generated during error-prone replication but not assembled into virus particles.

Interestingly, among the 21 isolates we sequenced, RNA3A was never found without RNA3B and vice versa. This observation aligns with previously sequenced HPWMoV isolates as well – in every sample where RNA3A or RNA3B were identified, another variant of RNA3 was present, even if it could not definitely be assigned to RNA3A or RNA3B. Seemingly, both RNA3A and RNA3B are required for type A isolates, yet type B isolates only require one version of RNA3, namely RNA3C. This suggests that RNA3C can somehow perform the combined function of RNA3A and RNA3B. It is not immediately apparent which RNA3 configuration is ancestral. A potential explanation would be that RNA3C is a recombinant of RNA3A and RNA3B, yet no evidence of recombination within RNA3C or any RNA3 sequences was found in this study. Previous studies have found predicted recombination events in RNA3 (Jones et al., 2023), but the authors did not specify if these were RNA3A/RNA3B recombinants. There is a greater risk of chimeric assembly of RNA3A and RNA3B sequences from short reads since both variants are always present within a sample, emphasizing the need for long-read sequencing to confirm predicted recombination events identified through assembled short reads, such as the recombinants predicted in the previous study. Alternatively, RNA3C may be ancestral, with RNA3A and RNA3B having evolved from two copies of RNA3C that then sufficiently specialized and diverged in function such that both variants are now required. In this study, we did find examples of duplicated RNA segments of the same variant, such as two versions of RNA6B in sample 23WA6, even though only one variant of RNA6 was sufficient for all other isolates. However, it is unclear what would drive the divergence of RNA3A and RNA3B. RNA3 encodes the nucleoprotein which encapsidates and protects the viral genome from degradation and would not be expected to rapidly evolve due to its crucial function. The reason for this heterogeneity in RNA3 is one of the biggest puzzles of HPWMoV biology. It has been hypothesized by others that P3-A might be needed for encapsidation of the virus in wheat and P3-B needed for encapsidation in the wheat curl mite, though it has not yet been definitively shown that HPWMoV replicates in its mite vector (Tatineni et al., 2014).

Varying numbers of variants seem to be tolerated for each genome segment, which is likely tied to the function of the resulting protein. RNA1 and RNA2 were often present as a single variant, and encode the viral RNA-dependent polymerase and envelope glycoprotein, respectively, and are thus crucial to virus replication and transmission and are, thus, less tolerant of divergence. RNA8, on the other hand, which can be present in as many as three variants, encodes one of two viral silencing suppressor proteins (Gupta et al., 2018), and as such, may be involved in a molecular arms race with the host antiviral immune system and, therefore, may be under more selective pressure to evolve. Virus isolates that maintain a more diverse complement of RNA8 variants may have a fitness advantage through the more rapid generation of novel RNA8 variants that can compete in the arms race. Curiously, the other silencing suppressor protein is encoded by RNA7, which, along with RNA1 and RNA2, was the least likely to be present as more than one variant in all isolates examined. It is possible that the tight evolutionary constraints on RNA7 are either the result of, and/or what allows, RNA8 to possess such variability. Studies into the mechanism of these two silencing suppressors suggest that they are distinct (Gupta et al., 2019). If reliable, basal silencing suppression is achieved by RNA7, then RNA8 is free to evolve to explore sequence space without a major fitness cost but with a potential fitness reward if a more favorable RNA8 variant develops. The contrasting evolutionary pressures on the two silencing suppressor proteins may also be related to their mechanisms; RNA7 may target a very conserved process in the host whereas RNA8 may interface with a more dynamic host defense process that requires rapid evolution. RNA8 was preliminarily found to bind dsRNA (Gupta et al., 2019), but the purified, assembled form of the P8 protein did not (Hamo et al., 2024), suggesting a yet-to-be-determined or even novel mechanism of silencing suppression.

Despite the variation tolerated on the individual genome segment level for some RNAs, overall there was no evidence of reassortment between type A and type B isolates in this study even though opportunities to reassort seem to occur, as evidenced by the two co-infected samples detected in this study. Such co-infections may occur quite readily as they were found twice in our limited sample size (9.5% of samples). The two isolate types are not geographically isolated either, as both types were readily found in the Columbia Basin. Despite the spurious addition of the alternate variant of some RNAs, such as RNA5B being present in the otherwise type A isolates 15WA, 22OH1, and 23OR4 in addition to the expected RNA5A, there were no examples of the alternate variant completely replacing the native variant in this study. Among previously sequenced isolates, there were four instances in which RNA5B seemed to replace RNA5A in the otherwise type A isolates: BCHPV1, BCHPV2, HP1G, and HPWMoV_NWB1. Considering that RNA5B was thrice found in type A isolates in this study, in addition to RNA5A, it is unclear if RNA5A was not present in these previously sequenced samples or just not detected as they were sequenced at a lower depth and with no virus-like particle enrichment. For instance, an RNA1 sequence was not recovered from sample HP1G. But the fact that this same phenomenon was observed numerous times across two very distant geographic regions (Australia and North America) suggested a true reassortment event, which would be the first evidence of reassortment for any HPWMoV genome segment. This general lack of reassortment hints that there is tight co-evolution among the RNA segments within a type and they are not generally interchangeable with the variants found in the other type (i.e., RNA1A is not interchangeable with RNA1B, etc.). This lack of reassortment explains why both variants of every RNA were preserved in the co-infected samples. This general lack of reassortment also represents a form of reproductive isolation between the two isolate types, which may drive further divergence. In fact, some historic reproductive isolation event is probably what gave rise to the two types in the first place – they would not have diverged if genetic exchange were still occurring. Though they share a common ancestor, it is possible that, eventually, type A and type B isolates will diverge enough to be considered separate emaravirus species infecting the same host. The species demarcation criterion for emaraviruses is that the gene products of two of the first four genome segments must diverge by greater than 20% on the amino acid level (Digiaro et al., 2024). While this is true of the gene products of RNA1A and RNA1B, RNAs 2, 3, and 4 differ by <20% on the amino acid level between types so they would not yet be considered separate viral species.

The geographic distribution of HPWMoV isolate types in sweet corn samples across the Pacific Northwest was quite striking, with both types as well as co-infections found in the Columbia Basin, but only type A isolates found in the Treasure Valley samples. Indeed, the previously sequenced HPWMoV isolate from Idaho was also type A. It is unclear what factors drive the diversity of isolate types in the Columbia Basin or the lack of diversity in the Treasure Valley. Notably, the Columbia Basin encompasses large-scale wheat production in addition to sweet corn production. Among our samples and others that have been sequenced to date, no clear relationship exists between isolate type and the plant host of collection as both isolate types have been found in corn and wheat. But if type B isolates originally came from wheat, or some other grass species present in the Columbia Basin but not the Treasure Valley, that could explain the lack of type B isolates in the Treasure Valley. HPWMoV may be using wheat or other winter grasses for overwintering, and while it has been shown that both isolate types infect wheat, it may be that the HPWMoV overwintering host(s) in the Treasure Valley can only support type A isolates. This uneven distribution of isolate types also suggests there is little genetic exchange between HPWMoV in the Columbia Basin and the Treasure Valley, or there is primarily one-way exchange from the Treasure Valley to the Columbia Basin, but not the inverse. It is also possible that type B isolates are present in the Treasure Valley but were not detected in this study as fewer samples were collected from this region relative to the Columbia Basin (7 vs. 13 samples).

The presence of both A and B type isolates in the Columbia Basin, a major area of sweet corn seed production and processing sweet corn production, does support the need to develop diagnostic assays that can detect both isolate types. In this study, we report two primer and probe sets that target two different RNA segments (RNA2 and RNA4) and can recognize both isolate types. All 20 isolates tested from this study were detected by both primer and probe sets, and *in silico* analysis of all previously sequenced HPWMoV isolates indicates that all but one would be detected by at least one of the primer and probe sets. Targeting these two RNA segments has the added benefit of reducing the chance of complication from multiple target RNA variants within the same sample, as RNA2 and RNA4 were often present as a single variant. Both primer sets also detected HPWMoV in the two co-infected samples tested in this study, and were effective on both leaf and seed tissue. Moreover, the two primer and probe sets also work as a duplex reaction for more efficient testing. The duplex reaction can be multiplexed further with a host endogenous control gene, if desired.

## 5. Conclusion

Overall, this study more than tripled the number of full-length HPWMoV genomes available and provides a new framework for classification of HPWMoV isolates into type A or type B. This study also advances our understanding of emaravirus genomic diversity and raises some interesting questions regarding emaravirus evolution. We also leveraged these sequences to develop a new, robust diagnostic assay for HPWMoV that is suitable for seed health testing for phytosanitary certification. More reliable diagnostics and an improved understanding of HPWMoV sequence diversity will, no doubt, aid future studies aimed towards understanding the virus’ distribution, evolutionary dynamics, and modes of transmission.

## Data Availability Statement

All HPWMoV sequences generated in this study have been deposited to GenBank (see **Supplementary Table 5** for accession numbers). Alignments of HPWMoV sequences generated in this study as well as alignments of all previously sequenced HPWMoV isolates by genome segment have been included as **Supplementary Data.** Raw Oxford Nanopore sequence results of predicted recombination regions are also included as **Supplementary Data.**

## Author Contributions

Conceptualization, funding acquisition, project administration, resources, supervision: J.R.W, E.W.O, L.J.d.T.; Investigation, Methodology, Validation: all authors; Formal analysis: J.R.W., E.W.O, K.J.W, N.K.; Software: E.W.O.; Data Curation: J.R.W., E.W.O, K.J.W.; Visualization: J.R.W., E.W.O.; Writing – Original Draft: J.R.W.; Writing – Review & Editing: all authors.

## Acknowledgements

We would like acknowledge industry collaborators who collected and shipped leaf samples that made this study possible. We would like to thank the Genomics Shared Resource at The James Comprehensive Cancer Center at The Ohio State University for library preparation and sequencing. We would also like to thank Li-Fang Chen of Bayer Vegetables for assistance in determining primer and probe concentrations for multiplexing. This work was supported by USDA Agricultural Research Service base funding, project #5082-22000-002-000D and the Western Integrated Pest Management Center which is funded via the Crop Protection and Pest Management Program, project #2022-70006-38003 from the USDA National Institute of Food and Agriculture.

## Supplementary Information for

## Supplementary Tables

**Supplementary Table 1.**
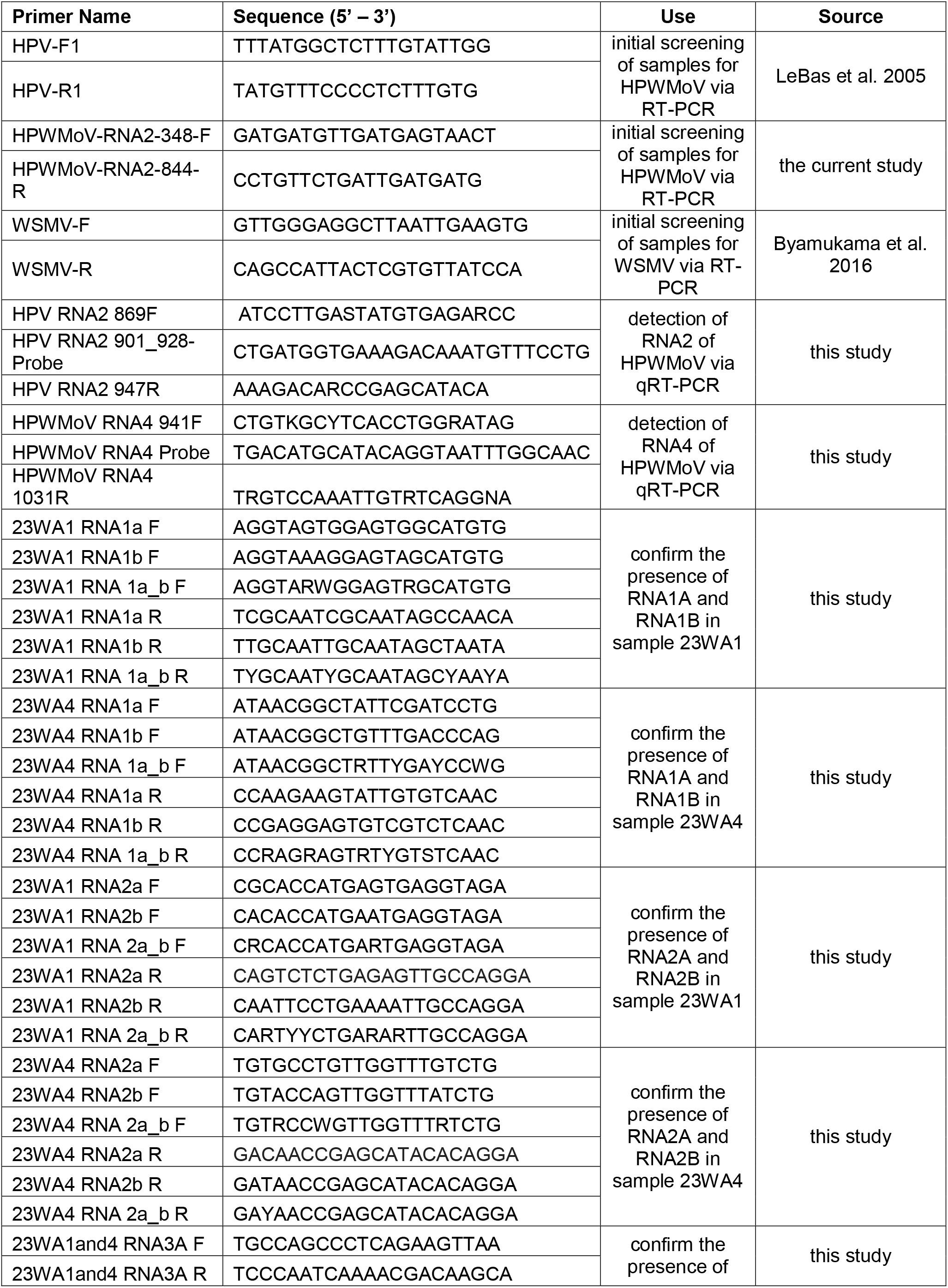
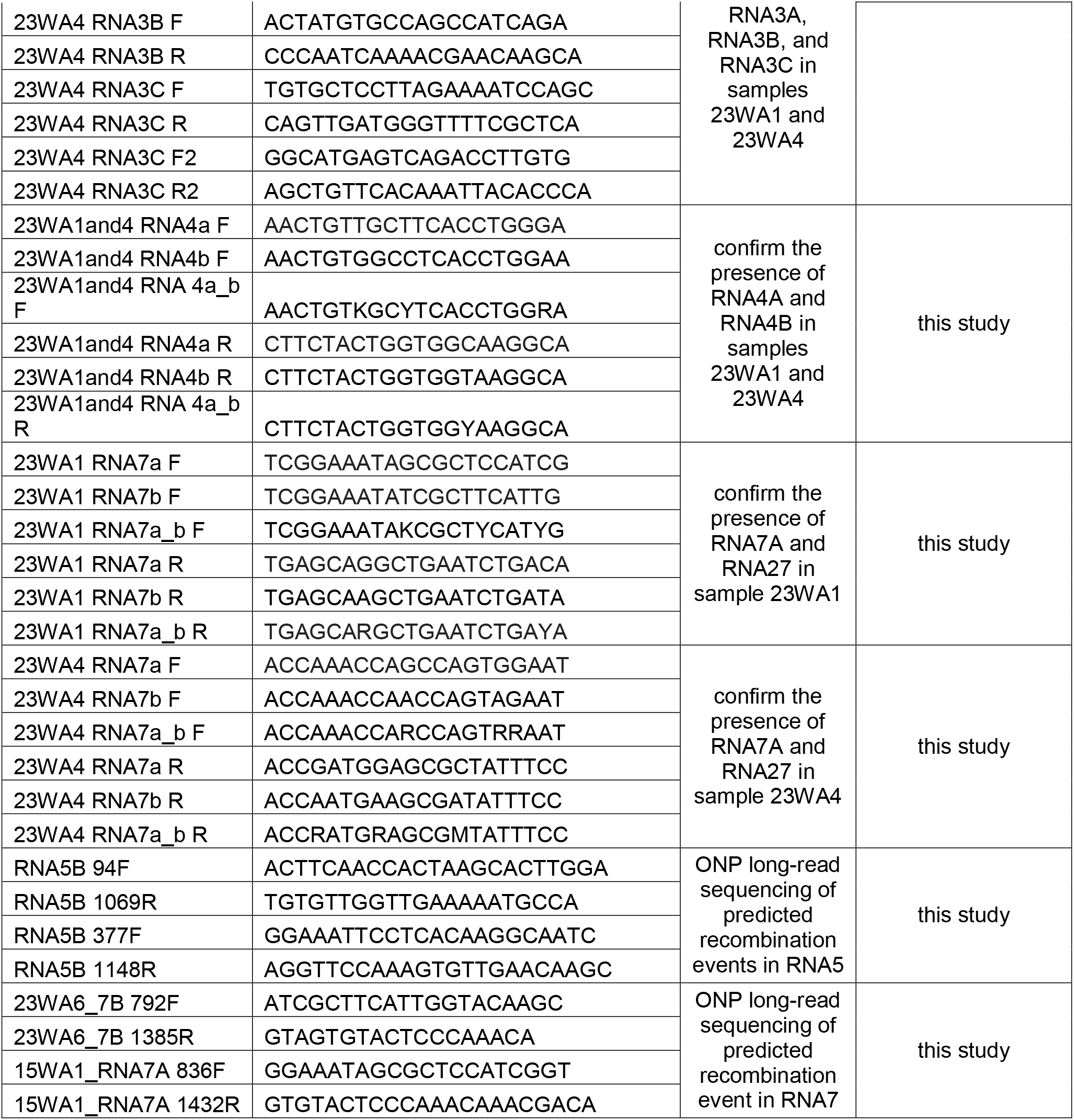
Primers and probes used in this study.

**Supplementary Table 2. Pairwise nucleotide percent identity between all variants of each RNA segment identified in this study and the reference sequence of HPWMoV

**Supplementary Table 3. Relative abundance of each RNA segment of HPWMoV within each sweet corn sample from the Pacific Northwest USA, presented as transcripts per million (TPM)

**Supplementary Table 4. Pairwise nucleotide percent identity between all previously sequenced variants of each RNA of HPWMoV and the RNA consensus sequences generated in this study

**Supplementary Table 5.**
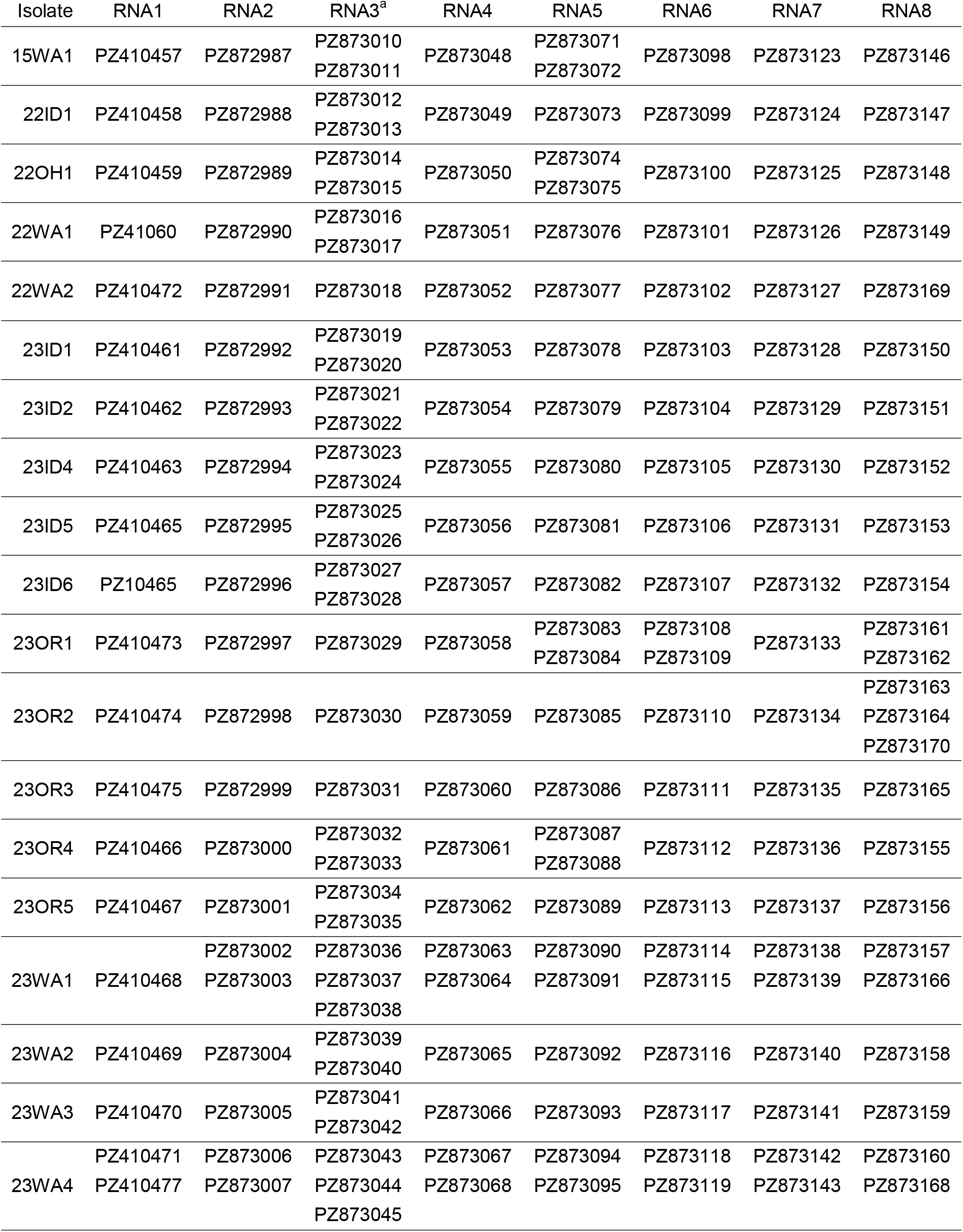
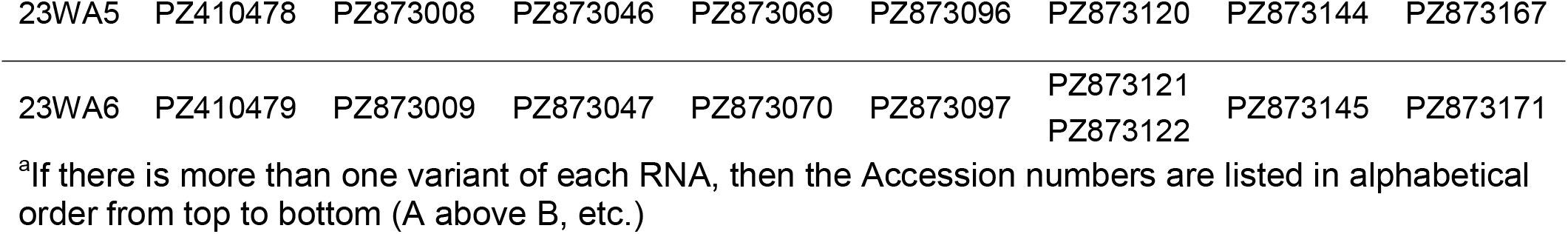
List of accession numbers of sequences of HPWMoV from this study deposited to GenBank.

**These tables are too large for this document and are attached as separate .xlsx files

## Supplementary Figures

**Supplemental Figure 1.**
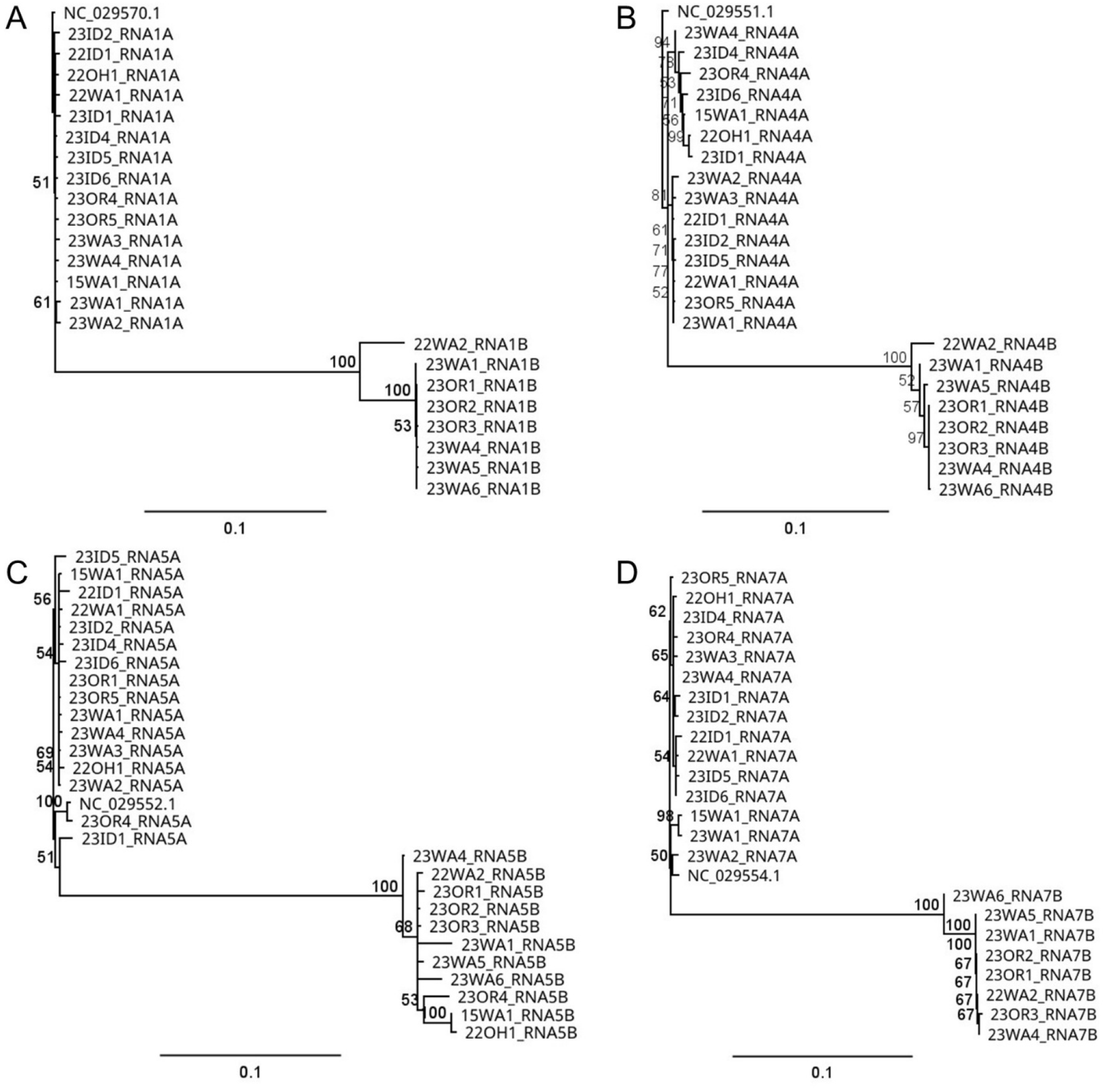
Phylogenetic trees for RNA1 (A), RNA4 (B), RNA5 (C), and RNA7 (D) of HPWMoV. Shown are phylogenetic trees for RNA1 (A), RNA4 (B), RNA5 (C), and RNA7 (D). The clade containing the reference sequence is given the designation “A” as in “RNA1A”, and the non-reference clade is designated “B” (i.e. “RNA1B”). Both panels are roughly to the same horizontal scale. Shown at the nodes are bootstrap values out of 100 replicates. Phylogenetic trees for the other RNA segments can be found in Figure 2 (RNA2, RNA3) and Figure 3 (RNA6, RNA8).

**Supplementary Figure 2.**
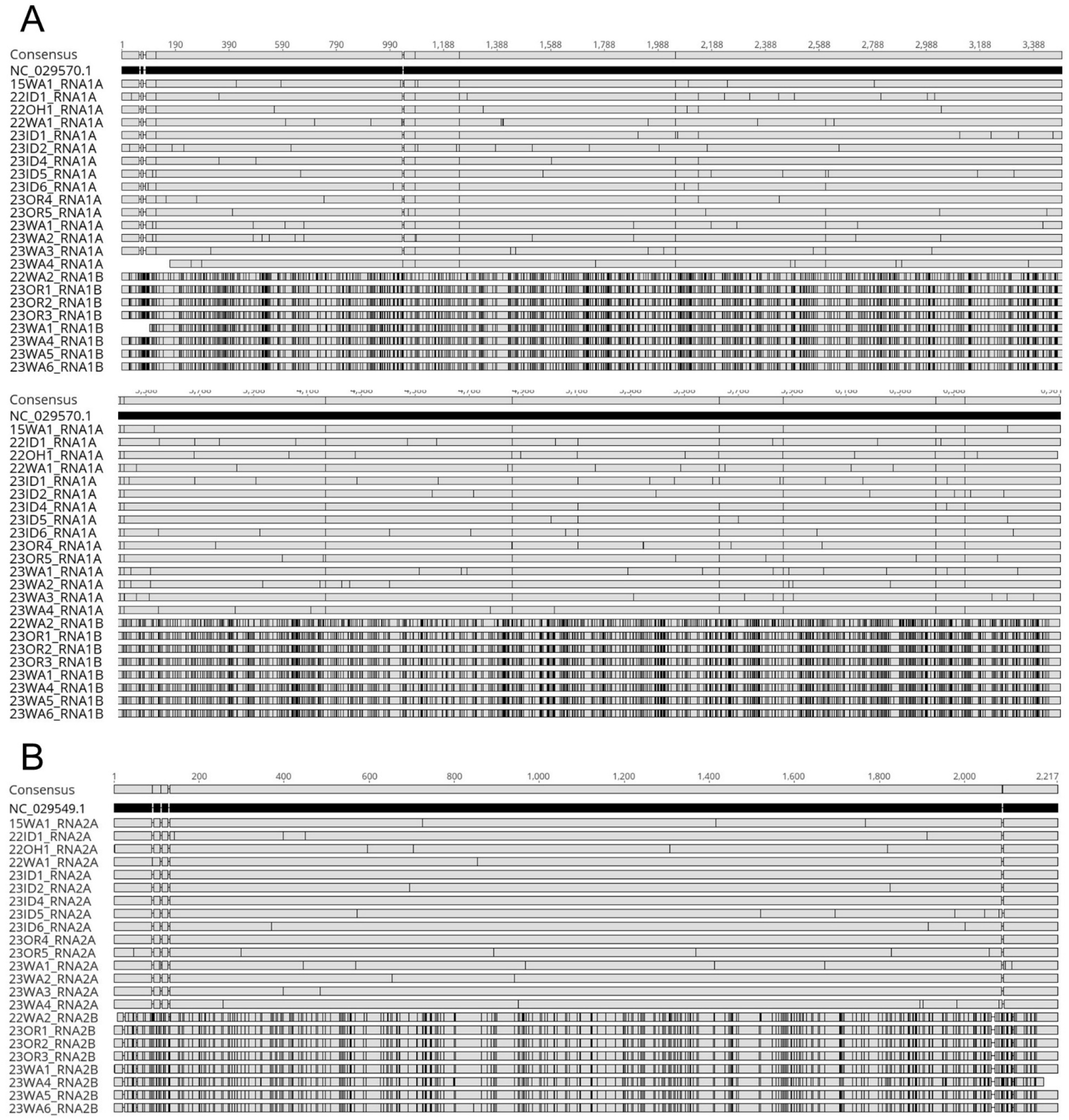
Multiple sequence alignment of all variants of RNA1 (A) and RNA2 (B) of HPWMoV identified in this study, aligned to the respective reference sequence. The reference sequence is shown as a black rectangle with all other sequences shown as gray rectangles (including the consensus sequence at the top, which includes the nucleotide number scale). Disagreement between an individual nucleotide in each sequence and the reference sequence is indicated with a black vertical line in the respective gray box. Gaps in an individual sequence appear as breaks in the respective rectangle, connected by a horizontal line.

**Supplementary Figure 3.**
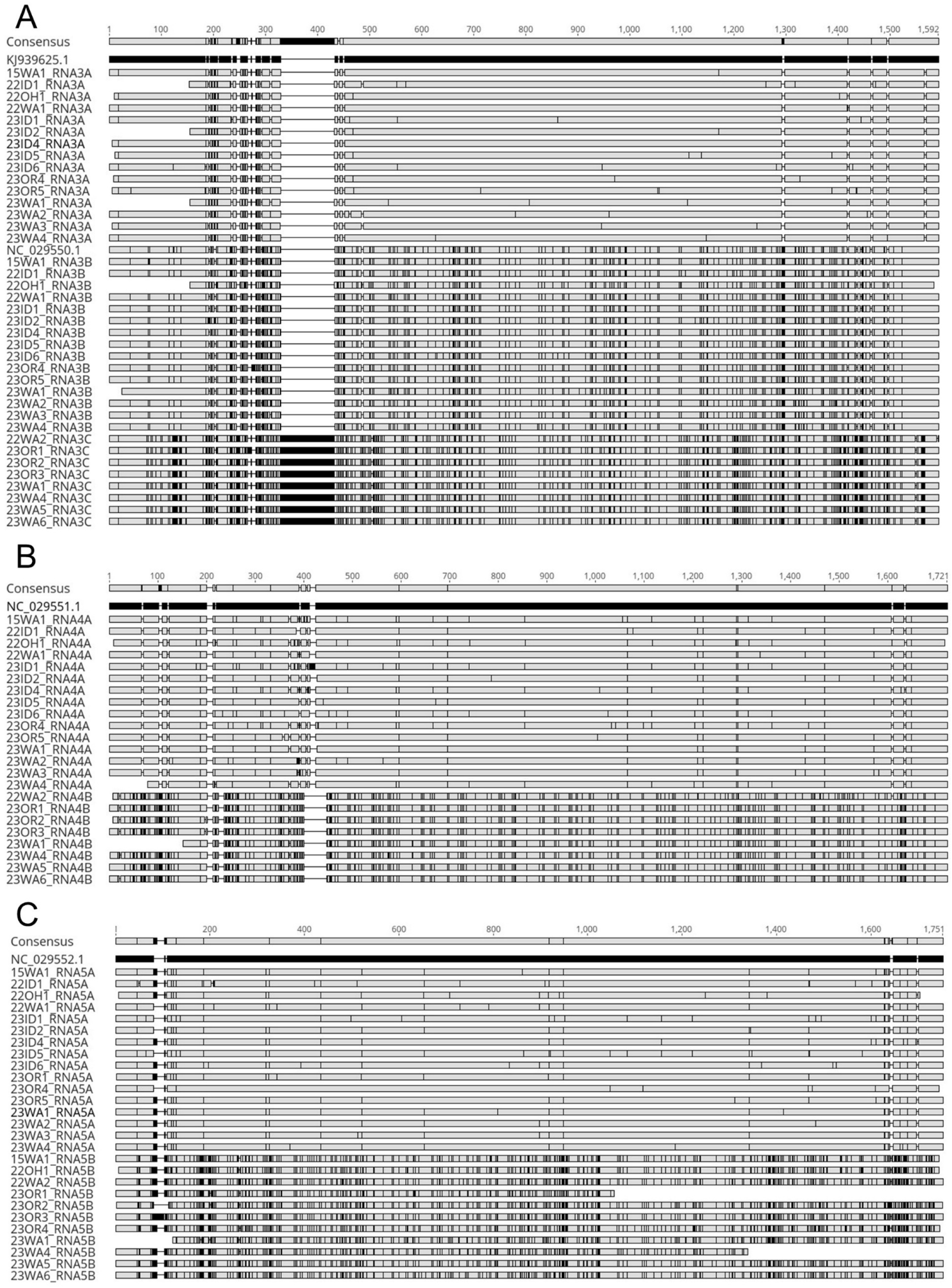
Multiple sequence alignment of all variants of RNA3 (A), RNA4 (B), and RNA5 (C) of HPWMoV identified in this study, aligned to the respective reference sequence. The reference sequence is shown as a black rectangle with all other sequences shown as gray rectangles (including the consensus sequence at the top, which includes the nucleotide number scale). Disagreement between an individual nucleotide in each sequence and the reference sequence is indicated with a black vertical line in the respective gray box. Gaps in an individual sequence appear as breaks in the respective rectangle, connected by a horizontal line.

**Supplementary Figure 4.**
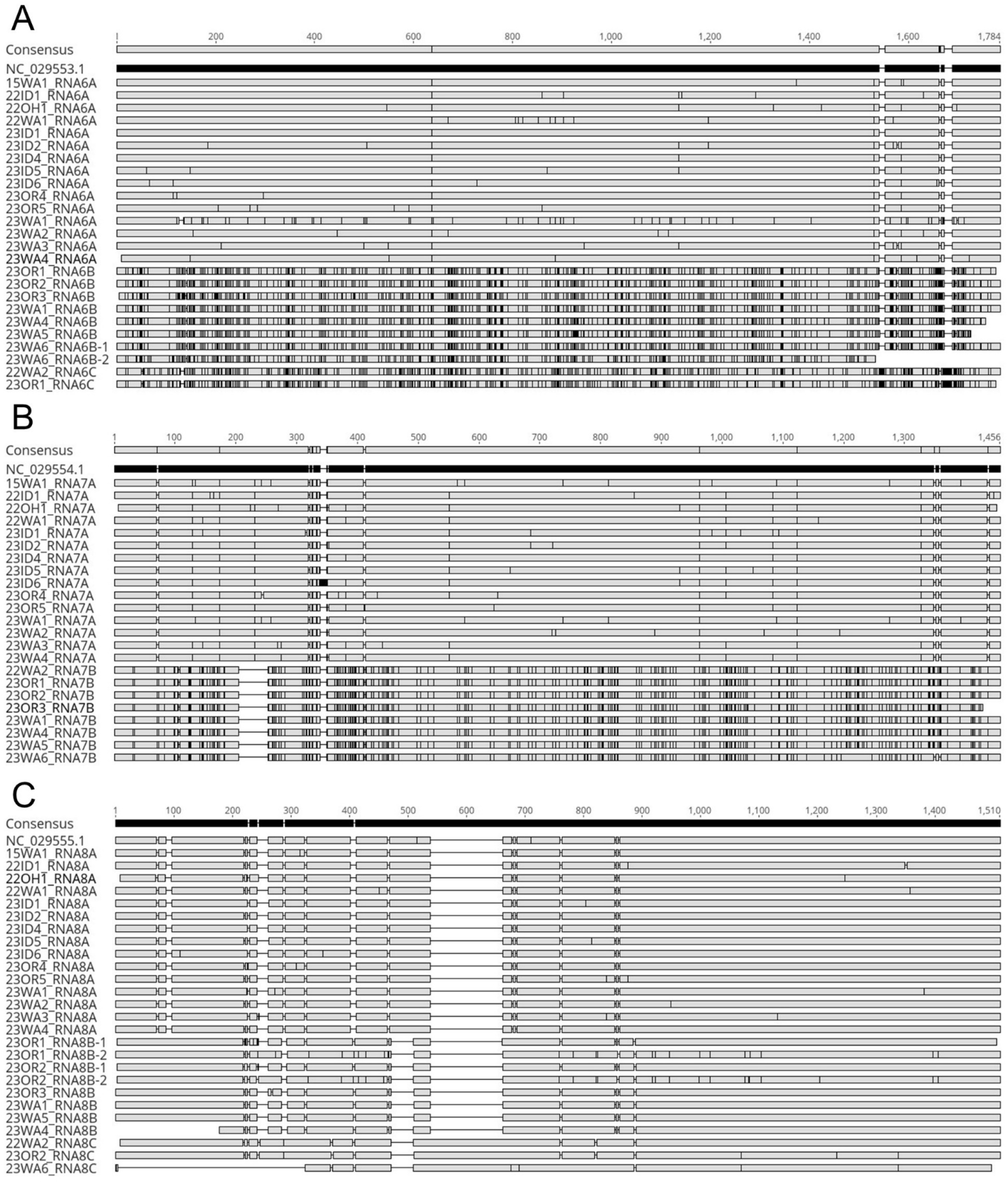
Multiple sequence alignment of all variants of RNA6 (A), RNA7 (B), and RNA8 (C) of HPWMoV identified in this study, aligned to the respective reference sequence. The reference sequence is shown as a black rectangle with all other sequences shown as gray rectangles (including the consensus sequence at the top, which includes the nucleotide number scale). Disagreement between an individual nucleotide in each sequence and the reference sequence is indicated with a black vertical line in the respective gray box. Gaps in an individual sequence appear as breaks in the respective rectangle, connected by a horizontal line.

**Supplementary Figure 5.**
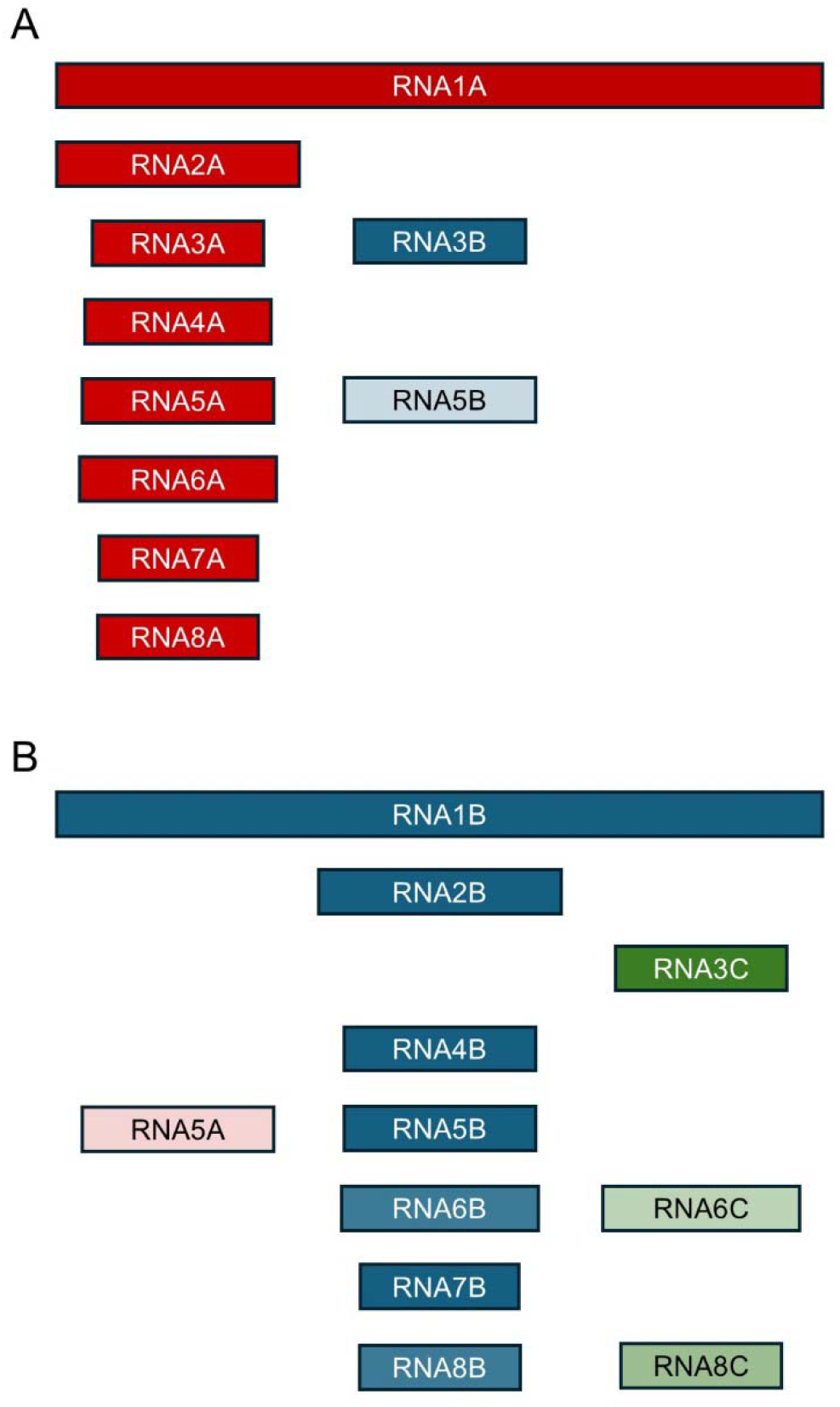
Visual representation of all variations of RNA segment composition of HPWMoV isolates sequenced in this study. Shown are variations of type A (A) and type B (B) isolates. Genome segments are represented by rectangles, which are to scale. “A” variants are red, “B” variants are blue, and “C” variants are green. The transparency of the coloring of each segment represents its relative appearance in that isolate type. For instance, RNA3B is always present in type A isolates whereas RNA5B was only present in 3 of 13 type A isolates in the current study and RNA5A was only found in 1 type B isolate

**Supplementary Figure 6.**
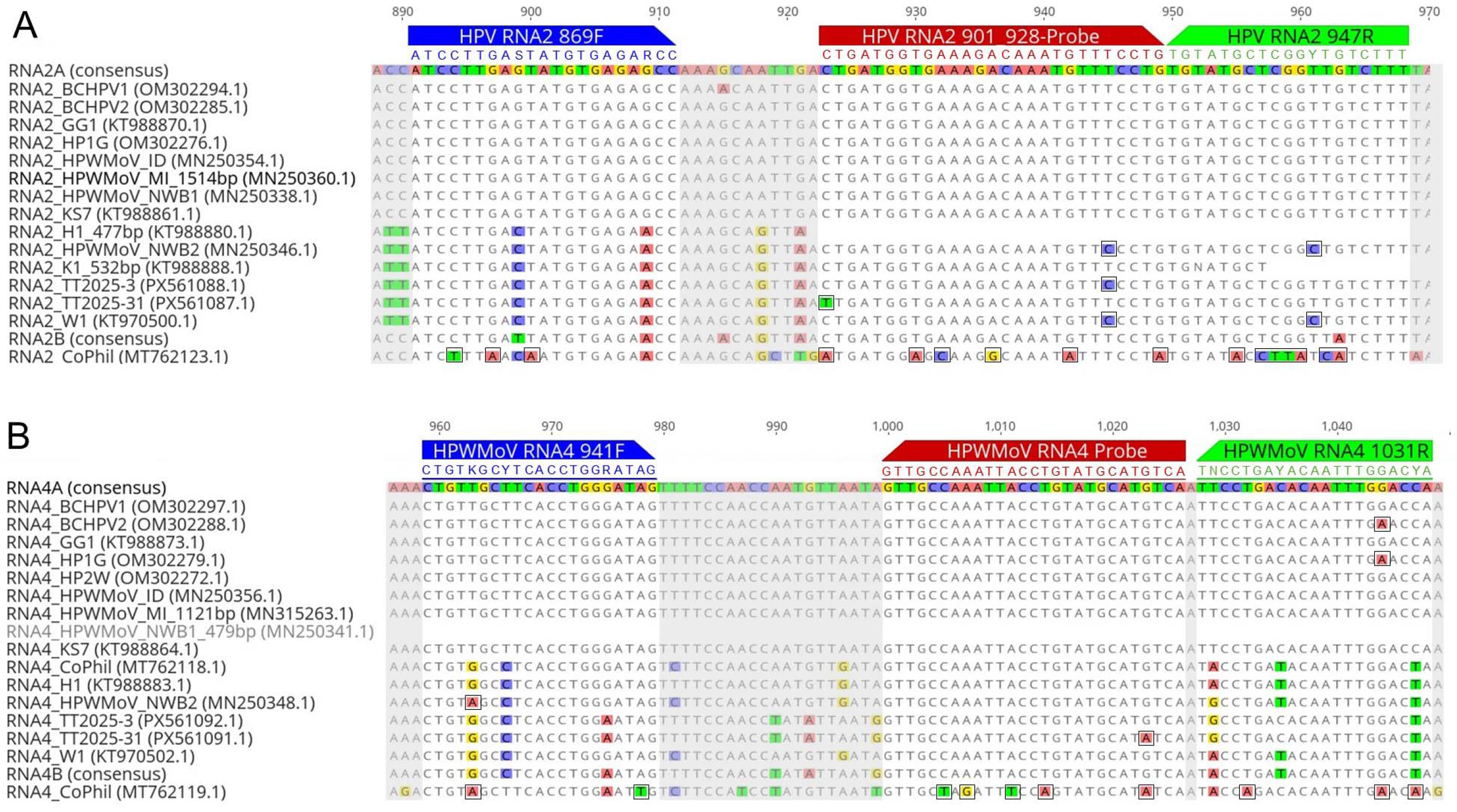
*In silico* prediction of RNA2 and RNA4 primer and probe set annealing to previously sequenced HPWMoV isolates. Pictured are the multiple sequence alignments for RNA2 and RNA4 in the region where the primer and probe sets anneal. Forward primers are shown in blue, reverse primers in green, and probes in red. Nucleotide differences from the consensus are highlighted. Differences that do not match the primer or probe sequences are outlined with a black box. Nucleotides not in the region where primers or probes anneal have been grayed out. The sequences for the reverse primer and the RNA4 probe are reverse complements so that mismatches could be more easily visibly identified as only the forward strand of DNA is presented here.

## Supplementary Data Files

Multiple sequence alignment of all RNA segments sequences generated in this study, aligned to the reference sequence of HPWMoV:

RNA1_CURRENT_ALN.fasta

RNA2_CURRENT_ALN.fasta

RNA3_CURRENT_ALN.fasta

RNA4_CURRENT_ALN.fasta

RNA5_CURRENT_ALN.fasta

RNA6_CURRENT_ALN.fasta

RNA7_CURRENT_ALN.fasta

RNA8_CURRENT_ALN.fasta

Multiple sequence alignment of all previously sequenced HPWMoV samples, aligned to the consensus sequences generated in this study:

RNA1_PREVIOUS_ALN.fasta

RNA2_PREVIOUS_ALN.fasta

RNA3_PREVIOUS_ALN.fasta

RNA4_PREVIOUS_ALN.fasta

RNA5_PREVIOUS_ALN.fasta

RNA6_PREVIOUS_ALN.fasta

RNA7_PREVIOUS_ALN.fasta

RNA8_PREVIOUS_ALN.fasta

Oxford Nanopore long-read sequencing results for predicted recombination events among RNA segments of HPWMoV sequenced in this study:

23WA5_RNA5B_region1.ab1

23WA5_RNA5B_region2.ab1

23WA6_RNA5B_region1.ab1

23WA6_RNA5B_region2.ab1

23WA6_RNA7B.ab1

15WA1_RNA7A.ab1

## References

Adegbola, R. O., Maheepala, D. C., Schuch, U. K., & Brown, J. K. (2025). Prevalence, host range, and characterization of multiple Palo verde broom emaravirus genomes and eriophyid mites from Parkinsonia spp. in Arizona. Virus Research, 361, 199643. 10.1016/j.virusres.2025.199643

Albrecht, T., White, S., Layton, M., Stenglein, M., Haley, S., & Nachappa, P. (2022). Occurrence of Wheat Curl Mite and Mite-Vectored Viruses of Wheat in Colorado and Insights into the Wheat Virome. Plant Disease, 106(10), 2678–2688. 10.1094/PDIS-02-21-0352-RE

Andrews, S. (2010). FastQC: a quality control tool for high throughput sequence data. In https://www.bioinformatics.babraham.ac.uk/projects/fastqc/

Arif, M., Aguilar-Moreno, G. S., Wayadande, A., Fletcher, J., & Ochoa-Corona, F. M. (2014). Primer modification improves rapid and sensitive in vitro and field-deployable assays for detection of high plains virus variants. Applied and Environmental Microbiology, 80(1), 320–327. 10.1128/AEM.02340-13

Bolger, A. M., Lohse, M., & Usadel, B. (2014). Trimmomatic: a flexible trimmer for Illumina sequence data. Bioinformatics, 30(15), 2114–2120. 10.1093/bioinformatics/btu170

Byamukama, E., Tatineni, S., Hein, G., McMechan, J., & Wegulo, S. N. (2016). Incidence of Wheat streak mosaic virus, Triticum mosaic virus, and Wheat mosaic virus in Wheat Curl Mites Recovered from Maturing Winter Wheat Spikes. Plant Disease, 100(2), 318–323. 10.1094/PDIS-06-15-0692-RE

Camacho, C., Coulouris, G., Avagyan, V., Ma, N., Papadopoulos, J., Bealer, K., & Madden, T. L. (2009). BLAST+: architecture and applications. BMC Bioinformatics, 10, 421. 10.1186/1471-2105-10-421

Candresse, T., Svanella-Dumas, L., Huang, A., Faure, C., Comte, R., & Marais, A. (2026). Characterization of French isolates of wheat mosaic virus and identification of multiple variants of genomic RNAs 5 and 6. Archives of Virology, 171(2), 35. 10.1007/s00705-025-06507-y

Digiaro, M., Elbeaino, T., Kubota, K., Ochoa-Corona, F. M., & von Bargen, S. (2024). ICTV Virus Taxonomy Profile: Fimoviridae 2024. Journal of General Virology, 105(5). 10.1099/jgv.0.001943

Establishes emergency measure to prevent the entry of High Plains virus (HPV) and wheat streak mosaic virus (WSMV) in corn seed from all origins. (6904/2022). (25 Nov 2022). Santiago, Chile: Gobierno de Chile, Servicio Agricola y Ganadero

Forster, R. L., Seifers, D. L., Strausbaugh, C. A., Jensen, S. G., Ball, E. M., & Harvey, T. L. (2001). Seed Transmission of the High Plains virus in Sweet Corn. Plant Disease, 85(7), 696–699. 10.1094/pdis.2001.85.7.696

Gibbs, M. J., Armstrong, J. S., & Gibbs, A. J. (2000). Sister-scanning: a Monte Carlo procedure for assessing signals in recombinant sequences. Bioinformatics, 16(7), 573–582. 10.1093/bioinformatics/16.7.573

Gupta, A. K., Hein, G. L., Graybosch, R. A., & Tatineni, S. (2018). Octapartite negative-sense RNA genome of High Plains wheat mosaic virus encodes two suppressors of RNA silencing. Virology, 518, 152–162. 10.1016/j.virol.2018.02.013

Gupta, A. K., Hein, G. L., & Tatineni, S. (2019). P7 and P8 proteins of High Plains wheat mosaic virus, a negative-strand RNA virus, employ distinct mechanisms of RNA silencing suppression. Virology, 535, 20–31. 10.1016/j.virol.2019.06.011

Hamo, S., Izhaki-Tavor, L. S., Tatineni, S., & Dessau, M. (2024). The RNA Silencing Suppressor P8 From High Plains Wheat Mosaic Virus is a Functional Tetramer. Journal of Molecular Biology, 436(24), 168870. 10.1016/j.jmb.2024.168870

Hodge, B. A., Paul, P. A., & Stewart, L. R. (2020). Occurrence and High-Throughput Sequencing of Viruses in Ohio Wheat. Plant Disease, 104(6), 1789–1800. 10.1094/pdis-08-19-1724-re

Holmes, E. C., Worobey, M., & Rambaut, A. (1999). Phylogenetic evidence for recombination in dengue virus. Molecular Biology and Evolution, 16(3), 405–409. 10.1093/oxfordjournals.molbev.a026121

Import Health Standard: Seeds for Sowing. (155.02.05). (Aug 30 2024). Wellington, New Zealand: New Zealand Government, Ministry for Primary Industries

Jones, R. A. C., Vazquez-Iglesias, I., McGreig, S., Fox, A., & Gibbs, A. J. (2023). Genomic High Plains Wheat Mosaic Virus Sequences from Australia: Their Phylogenetics and Evidence for Emaravirus Recombination and Reassortment. Viruses, 15(2). 10.3390/v15020401

Lam, H. M., Ratmann, O., & Boni, M. F. (2018). Improved Algorithmic Complexity for the 3SEQ Recombination Detection Algorithm. Molecular Biology and Evolution, 35(1), 247–251. 10.1093/molbev/msx263

Lebas, B. S. M., Ochoa-Corona, F. M., Elliott, D. R., Tang, Z., & Alexander, B. J. R. (2005). Development of an RT-PCR for High Plains virus Indexing Scheme in New Zealand Post-Entry Quarantine. Plant Disease, 89(10), 1103–1108. 10.1094/pd-89-1103

Li, H. (2018). Minimap2: pairwise alignment for nucleotide sequences. Bioinformatics, 34(18), 3094–3100. 10.1093/bioinformatics/bty191

Li, H., & Durbin, R. (2009). Fast and accurate short read alignment with Burrows-Wheeler transform. Bioinformatics, 25(14), 1754–1760. 10.1093/bioinformatics/btp324

Louie, R., Seifers, D. L., & Bradfute, O. E. (2006). Isolation, transmission and purification of the High Plains virus. Journal of Virological Methods, 135(2), 214–222.

Martin, D., & Rybicki, E. (2000). RDP: detection of recombination amongst aligned sequences. Bioinformatics, 16(6), 562–563. 10.1093/bioinformatics/16.6.562

Martin, D. P. (2020). RDP5 Instruction Manual. 1-52. Retrieved July 2026, from https://web.cbio.uct.ac.za/~darren/RDP5Manual.pdf

Martin, D. P., Posada, D., Crandall, K. A., & Williamson, C. (2005). A modified bootscan algorithm for automated identification of recombinant sequences and recombination breakpoints. AIDS Research and Human Retroviruses, 21(1), 98–102. 10.1089/aid.2005.21.98

Martin, D. P., Varsani, A., Roumagnac, P., Botha, G., Maslamoney, S., Schwab, T., Kelz, Z., Kumar, V., & Murrell, B. (2021). RDP5: a computer program for analyzing recombination in, and removing signals of recombination from, nucleotide sequence datasets. Virus Evol, 7(1), veaa087. 10.1093/ve/veaa087

Nischwitz, C. (2020). Seed-Transmitted Wheat Mosaic Virus in Sweet Corn in Utah. Plant Health Progress, 21(3), 212–213. 10.1094/php-12-19-0092-br

Padidam, M., Sawyer, S., & Fauquet, C. M. (1999). Possible emergence of new geminiviruses by frequent recombination. Virology, 265(2), 218–225. 10.1006/viro.1999.0056

Patro, R., Duggal, G., Love, M. I., Irizarry, R. A., & Kingsford, C. (2017). Salmon provides fast and bias-aware quantification of transcript expression. Nat Methods, 14(4), 417–419. 10.1038/nmeth.4197

Posada, D., & Crandall, K. A. (2001). Evaluation of methods for detecting recombination from DNA sequences: computer simulations. Proceedings of the National Academy of Sciences of the United States of America, 98(24), 13757–13762. 10.1073/pnas.241370698

Pozhylov, I., Snihur, H., Shevchenko, T., Budzanivska, I., Liu, W., Wang, X., & Shevchenko, O. (2022). Occurrence and Characterization of Wheat Streak Mosaic Virus Found in Mono- and Mixed Infection with High Plains Wheat Mosaic Virus in Winter Wheat in Ukraine. Viruses, 14(6). 10.3390/v14061220

Prjibelski, A., Antipov, D., Meleshko, D., Lapidus, A., & Korobeynikov, A. (2020). Using SPAdes De Novo Assembler. Curr Protoc Bioinformatics, 70(1), e102. 10.1002/cpbi.102

Seifers, D. L., Harvey, T. L., Martin, T. J., & Jensen, S. G. (1998). A Partial Host Range of the High Plains Virus of Corn and Wheat. Plant Disease, 82(8), 875–879. 10.1094/PDIS.1998.82.8.875

Smith, J. M. (1992). Analyzing the mosaic structure of genes. Journal of Molecular Evolution, 34(2), 126–129. 10.1007/BF00182389

Stewart, L. R. (2016). Sequence diversity of wheat mosaic virus isolates. Virus Research, 213, 299–303. 10.1016/j.virusres.2015.11.013

Tatineni, S., McMechan, A. J., Wosula, E. N., Wegulo, S. N., Graybosch, R. A., French, R., & Hein, G. L. (2014). An eriophyid mite-transmitted plant virus contains eight genomic RNA segments with unusual heterogeneity in the nucleocapsid protein. Journal of Virology, 88(20), 11834–11845.

Tatineni, S., Ziems, A. D., Wegulo, S. N., & French, R. (2009). Triticum mosaic virus: a distinct member of the family potyviridae with an unusually long leader sequence. Phytopathology, 99(8), 943–950. 10.1094/PHYTO-99-8-0943

Weiller, G. F. (1998). Phylogenetic profiles: a graphical method for detecting genetic recombinations in homologous sequences. Molecular Biology and Evolution, 15(3), 326–335. 10.1093/oxfordjournals.molbev.a025929

Xie, W., Marty, D. M., Xu, J., Khatri, N., Willie, K., Moraes, W. B., & Stewart, L. R. (2021). Simultaneous gene expression and multi-gene silencing in Zea mays using maize dwarf mosaic virus. BMC Plant Biology, 21(1), 208. 10.1186/s12870-021-02971-1

